# True time series of gene expression from multinucleate single cells reveal essential information on the regulatory dynamics of cell differentiation

**DOI:** 10.1101/2020.09.16.299578

**Authors:** Anna Pretschner, Sophie Pabel, Markus Haas, Monika Heiner, Wolfgang Marwan

## Abstract

Dynamics of cell fate decisions are commonly investigated by inferring temporal sequences of gene expression states by assembling snapshots of individual cells where each cell is measured once. Ordering cells according to minimal differences in expression patterns and assuming that differentiation occurs by a sequence of irreversible steps, yields unidirectional, eventually branching Markov chains with a single source node. In an alternative approach, we used multinucleate cells to follow gene expression taking true time series. Assembling state machines, each made from single-cell trajectories, gives a network of highly structured Markov chains of states with different source and sink nodes including cycles, revealing essential information on the dynamics of regulatory events. We argue that the obtained networks depict aspects of the Waddington landscape of cell differentiation and characterize them as reachability graphs that provide the basis for the reconstruction of the underlying gene regulatory network.

## Introduction

Single-cell analyses revealed complex dynamics of gene regulation in differentiating cells (Junker and van Oudenaarden 2014; Marr et al. 2016; Paul et al. 2015; Plass et al. 2018; Spiller et al. 2010). It is believed that dynamic effects possibly superimposed by stochastic fluctuations in gene expression levels may play crucial roles in cell fate choice, commitment, and reprogramming (Bornholdt and Kauffman 2019; Ferrell Jr 2012; Graf and Enver 2009; Huang et al. 2009; Il Joo et al. 2018; Zhou and Huang 2011). Changes in gene expression over time have not been directly measured in single mammalian cells as cells are – for technical reasons – sacrificed during the analysis procedure and hence can be measured only once. Instead, algorithms have been developed to infer the gene expression trajectory of a typical cell in pseudo-time from static snapshots of gene expression states in a cell population, resulting in Markov chains of states (Bendall et al. 2014; Cannoodt et al. 2016; Chen et al. 2019; Saelens et al. 2019; Setty et al. 2019). Most trajectory inference algorithms are based on the assumption that differentiation is unidirectional (Bendall et al. 2014; Haghverdi et al. 2016; Saelens et al. 2019; Street et al. 2018) and that the probability of transiting from one state to the next similar state is independent of the individual history of a cell (Setty et al. 2019). The inference of trajectories has been used to create pseudo-time series for differentiation (Macaulay et al. 2016; Marco et al. 2014; Moignard et al. 2015; Shin et al. 2015), cell cycle (Kafri et al. 2013), and the response to perturbation (Gaublomme et al. 2015). As any given distribution of expression patterns could result from multiple dynamics, the reconstruction of trajectories from snapshots faces fundamental limits (Weinreb et al. 2018). Even though regulatory mechanisms cannot be directly and rigorously inferred from snapshots (Weinreb et al. 2018), dynamic analyses may be of immediate importance to resolve competing views on basic mechanisms and the role of stochasticity in cell fate decisions (Moris et al. 2016).

True single cell time series can be obtained in *Physarum polycephalum* by taking multiple samples of one and the same giant cell. *Physarum* belongs to the amoebozoa group of organisms. It has a complex, prototypical eukaryote genome (Schaap et al. 2016) and forms different cell types during its life cycle (Alexopoulos and Mims 1979).

Giant, multi-nucleate cells, so-called plasmodia provide a source of macroscopic amounts of homogeneous protoplasm with a naturally synchronous population of nuclei, which is continually mixed by vigorous shuttle-streaming (Dove et al. 1986; Guttes and Guttes 1961, 1964; Rusch et al. 1966). The differentiation of a plasmodium into fruiting bodies involves extensive remodeling of signal transduction and transcription factor networks with alterations at the transcriptional, translational, and post-translational level (Glöckner and Marwan 2017).

In starving plasmodial cells, the formation of fruiting bodies can be experimentally triggered by a brief pulse of far-red light received by phytochrome as photoreceptor (Lamparter and Marwan 2001; Schaap et al. 2016; Starostzik and Marwan 1995b). Retrieving small samples of the same plasmodial cell before and at different time points after an inductive light pulse allows to follow how gene expression changes over real time. Because cell cycle, cell fate choice, and development are synchronous throughout the plasmodium (Hoffmann et al. 2012; Rätzel and Marwan 2015; Rusch et al. 1966; Starostzik and Marwan 1995a; Walter et al. 2013), single-cell gene expression trajectories can indeed be constructed from time series. By assembling finite state machines made from trajectories we have constructed Petri net models for the state transitions that predict Markov chains as variable developmental routes to differentiation (Rätzel et al. 2020; Werthmann and Marwan 2017) which may be considered as trajectories through the Waddington landscape (Huang et al. 2009; Waddington 1957). These Petri nets also predict reversible and irreversible steps, commitment points, and meta-stable states in cells responding to a differentiation stimulus. However, the computational approach for the construction of Petri nets from time series has been originally developed with data sets of a coarse resolution in time and the structural resolution of the nets was accordingly limited. Nevertheless, the approach turned out to be useful for capturing the dynamics of the process. For this paper, we developed a method for retrieving smaller samples from even larger plasmodial cells and showed that these cells provide a homogenious source for samples to be taken. This allowed us to considerably improve the time resolution as compared to previous studies. Sampling cells at higher time resolution, allowed the construction of Petri nets with enhanced structural and dynamic resolution. Structural complexity, highly connected nodes, parallel pathways, reversible reactions, and Petri net places representing meta-stable states in the developmental network, as revealed by the new data sets, characterize the differentiation response as complex and dynamic in contrast to a smooth, continuous process. We conclude that the dynamics revealed by our analysis most likely emerge from the non-linear dynamic behaviour of the underlying regulatory network rather than from stochastic fluctuations in the concentration of regulatory molecules.

## Results

### Even large plasmodia provide a source of homogeneous cell material for true time series analysis of gene expression

In previous studies we have shown that the gene expression pattern in samples taken at the same time from different sites of a plasmodium covering a standard Petri dish (9 cm Ø) did not change within the limits of accuracy of the measurements. Accordingly, repeated sampling of the same plasmodial cell yields true time series (Rätzel 2015; Werthmann and Marwan 2017). To allow more samples to be taken without consuming too much of the plasmodial mass, we now prepared plasmodia on 14 cm Ø Petri dishes, increasing the surface area covered by the plasmodial mass by 2.4-fold, and took smaller samples by punching agar plugs of 1.13 cm^2^ per sample, to harvest a small portion of the initial total plasmodial mass. To test whether the homogeneity in gene expression is impaired or even lost in the larger plasmodia, we took 9 or 16 samples at the same time from approximately evenly spread sites of a plasmodium, and estimated the gene expression pattern twice in the RNA of each sample, to obtain one technical replicate of each measurement (Aselmeyer 2019; Driesch 2019). This allowed to estimate the biological variation in gene expression within a plasmodium as compared to the technical accuracy of the measurements. In order to correct for potential differences in the efficiency of the RT-PCR between reactions, the expression value for each gene was normalized to the median of the expression values of all genes measured in the sample (for details see Materials and Methods). To estimate the technical accuracy of the measurements, the relative deviation of first and second measurement from the mean of the two measurements was estimated for each assayed gene in each of the retrieved plasmodial samples. The frequency distribution of all values was almost symmetrical with a tail consisting of a small number of low values, obviously as a result of inefficient RT-PCR reactions. To estimate the degree of homogeneity in gene expression within a plasmodial cell, we asked to which extent the expression values for the 9 or 16 samples taken from the same plasmodium deviated from their median. To restrict the influence of technical artefacts on the result, we considered the subset of the data where first and the second measurement of the same plasmodial sample deviated not more than two-fold from the mean of the two values. In three of the total of 46 analysed plasmodia (30 far-red stimulated; 16 dark controls), individual samples deviated from the rest of the samples of the same plasmodium by more than a factor of two. As errors in sample preparation could not be ruled out, these three plasmodia were excluded and the remaining data set of 43 plasmodia (28 far-red stimulated; 15 dark controls) was analysed taking the mean of 1st and 2nd measurement for each gene in each sample. Among the total of 7160 values, 98 % of the symmetric frequency distribution (SI Fig. 1) were between 0.48-fold and 2.10-fold deviation of the median of all values of the respective plasmodium (SI Table 1). There was no obvious difference between dark controls and far-red stimulated plasmodia which were measured at 6h after the light pulse when genes were already differentially regulated (SI Fig. 1; SI Table 1), indicating that even during the period where the mRNA abundance changed in time, the homogeneity in gene expression levels is maintained. Visual inspection of outliers within the distribution did not reveal any candidates for specific genes that might be inhomogeneously expressed.

In summary, the gene expression values throughout a plasmodium deviated not more than approximately two-fold from the median of all samples from the same plasmodium and were thus within the limits of the technical accuracy of the measurements, even under conditions were genes were in the process of being up- or down-regulated. These differences measured between samples were minor as compared to the differential regulation where the expression level of genes changed in the order of ten to more than hundred-fold (SI Fig. 2). These results are consistent with the results of the time-series experiment, where for each time point, two samples were retrieved and analysed from the same plasmodial cell (see below).

### Sampling of plasmodia at 1h time interval

As the assayed genes were evenly expressed and changed evenly in time throughout the large plasmodia, at least within the limits of accuracy of the measurements, we took time series at 1h time intervals. In order to assay, in each experiment, for the homogeneity and synchrony in gene expression throughout the plasmodium, we took two samples at each time point from different, arbitrarily chosen but distant sites of the plasmodium. In far-red stimulated plasmodia, the first samples (referred to as the 0h samples) were taken at the start of the experiment, *i.e*. immediately before application of the 15 min pulse of far-red light. All subsequent samples were taken at 1h time intervals until 10h after the start of the experiment. (At 5 to 6h after the far-red pulse cells have passed the commitment point, while visible morphogenesis starts several hours later by entering the transient nodulation stage at about 11h after the pulse (Hoffmann et al. 2012)). In the dark controls, the far-red stimulus was omitted. Gene expression in each plasmodial sample taken at a given time point was analysed twice by GeXP-RT-PCR, where the measurement and the corresponding technical replicate are referred to as 1st and 2nd measurement for sample #1, and 3rd and 4th measurement for sample #2, respectively. Data were normalized as described above.

The technical quality of the measurements was estimated separately for the two data sets, each comprising the data of the samples collected at the 11 time points of the time series. For each plasmodial sample, the relative deviation of the two measurements (1st and 2nd, or 3rd and 4th) from the mean of the two measurements was estimated. The frequency distributions of the deviations and corresponding quantil values indicated that the technical quality of 1st and 2nd, as well as 3rd and 4th measurement were virtually identical with 95% of the values differing less than a factor of two from each other (Fig. 1; Table 1).

**Table 1.** Quantil distributions of the x-fold deviation (x) of a value from the mean of two or four values, characterizing the reproducibility of measurements as estimated through technical and biological replicates, repectively. The table quantitatively characterizes the frequency distributions shown in Fig. 1.

| Measurements | 1st & 2nd |  | 3rd & 4th |  | 1st to 4th |  |
| --- | --- | --- | --- | --- | --- | --- |
| Percent of values | Quantile (Log2 (x)) | Quantile (x) | Quantile (Log2 (x)) | Quantile (x) | Quantile (Log2 (x)) | Quantile (x) |
| 1% | -1.310 | 0.403 | -1.379 | 0.384 | -1.346 | 0.393 |
| 5% | -0.555 | 0.681 | -0.592 | 0.663 | -0.573 | 0.672 |
| 25% | -0.167 | 0.890 | -0.169 | 0.890 | -0.168 | 0.890 |
| 50% | -0.010 | 0.993 | -0.008 | 0.994 | -0.009 | 0.994 |
| 75% | 0.137 | 1.100 | 0.141 | 1.102 | 0.139 | 1.101 |
| 95% | 0.415 | 1.333 | 0.415 | 1.333 | 0.415 | 1.333 |
| 99% | 0.751 | 1.683 | 0.747 | 1.679 | 0.750 | 1.682 |

**Figure 1.**
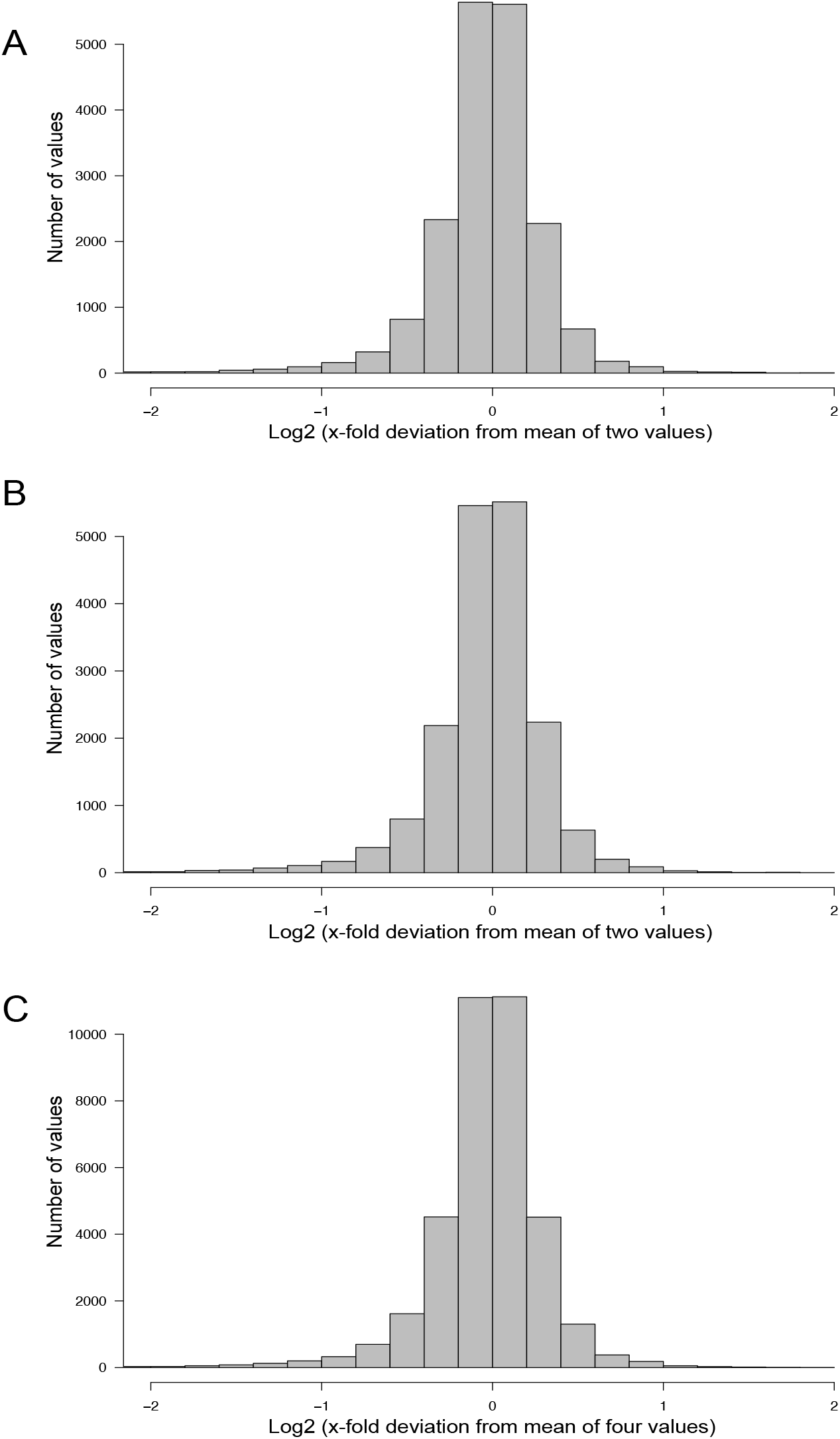
Technical accuracy of measurements and homogeneity of gene expression within a plasmodial cell as determined by technical and biological replicates, taken in experiments #1 and #2 (Table 2). (A, B) Technical accuracy of measurements of gene expression. The concentration of the mRNAs of the set of 35 genes (SI Table 2) was determined twice by RT-PCR for each RNA sample. The frequency distributions display the Log2 of the x-fold deviation of each expression value of each gene from the mean of the two values obtained by technical replication. Panels (A) and (B) show the results obtained for each of the two biological samples ((A), sample #1; (B), sample #2), that both were simultaneously taken from the same plasmodial cell at any time point during the experiments. C) Combination of the data sets shown in (A) and (B). This frequency distribution shows the deviation of each measurement from the mean of four values, obtained by twice measuring each of the two biological samples simultaneously taken from the same plasmodium at any time point of the experiments. The figure represents the complete data set of 36,540 data points that was analysed in the present study.

The degree of spatial variability of gene expression within a plasmodium was estimated by combining the data sets for the first and the second sample of a plasmodium taken at each time point of the time series. The frequency distribution of the deviation of each measurement from the mean of 1st, 2nd, 3rd, and 4th measurement of the two samples taken from each plasmodium at any time point was virtually identical to the frequency distributions obtained for the technical replicates, indicating that gene expression within the analysed plasmodia varied at maximum within the limits of accuracy of the measurements (within a factor of 2 in 95 % of the samples). This conclusion is based on the comparison of the quantile distributions of the data sets (Fig. 1; Table 1) considering a total of 36,540 data points.

### Multi-dimensional scaling analysis

With this data set, we investigated how expression changes as a function of time in the individual plasmodial cells. The gene expression pattern of a plasmodial cell at a given time point was obtained as the mean of the four expression values of each gene measured in the two plasmodial samples picked at that time point.

For visual representation of the data set and of single-cell trajectories of gene expression, we performed multidimensional scaling (MDS) to obtain a data point for the expression pattern of each cell at each time point. Single-cell trajectories of gene expression are shown in Fig. 2. Notably, the gene expression patterns of un-stimulated cells (dark controls) changed as a function of time with the highest variability along coordiate 2 of the MDS plot Fig. 2A,B. Trajectories of far-red stimulated cells (Fig. 2C,D) moved from the left side to the right side of the plot, while the shape of individual trajectories varied to a certain extent, indicating that the response of the cells was similar though not identical. Obviously, the trajectories of six of the eight far-red-stimulated plasmodia of experiment #1 traversed a considerably larger area of the MDS plot (Fig. 2C) as compared to the other stimulated cells, indicating a larger variation in gene expression during the response to the stimulus. The extent of variation is accordingly obvious when the bulk of data points is placed in the same plot (Fig. 3A). To search for genes that may account for the scattering along coordinate 2, we visually inspected the individual time series displayed in the form of a heat map (SI Fig 2). In addition to the genes that were clearly up- or down-regulated in response to the stimulus, the messages of four genes, *hstA, nhpA, pcnA,* and *uchA,* in the following called *pcnA-group* genes, changed over time in some of the plasmodia, but there was no obvious consistent relationship to the time point of stimulus application. When only genes were included in the analysis that were clearly up- or down-regulated in response to the stimulus (SI Fig 2, see also Fig. 7), the data points of the MDS plot were indeed less scattered (Fig 3B). A qualitatively similar result was obtained plotting the up-regulated and the down-regulated genes separately (Fig 3C,D), with some more variation in the expression of the down-regulated genes. However, expression of the *pcnA*-group over time (Fig 3E) was clearly different from the up- or down-regulated genes. Expression of the *pcnA*-group genes was different between cells from experiments #1 and #2 as seen from the trajectories of the cells (SI Fig 3), suggesting that ongoing internal processes in cells of experiment #1 might even influence their response to the light stimulus. Indeed, according to the corresponding MDS plots of the bulk data points (SI Fig 4) the response of the up- and down-regulated genes was less uniformly in cells of experiment #1 (SI Fig 4C) as compared to those of experiment #2 (SI Fig 4D).

**Figure 2.**
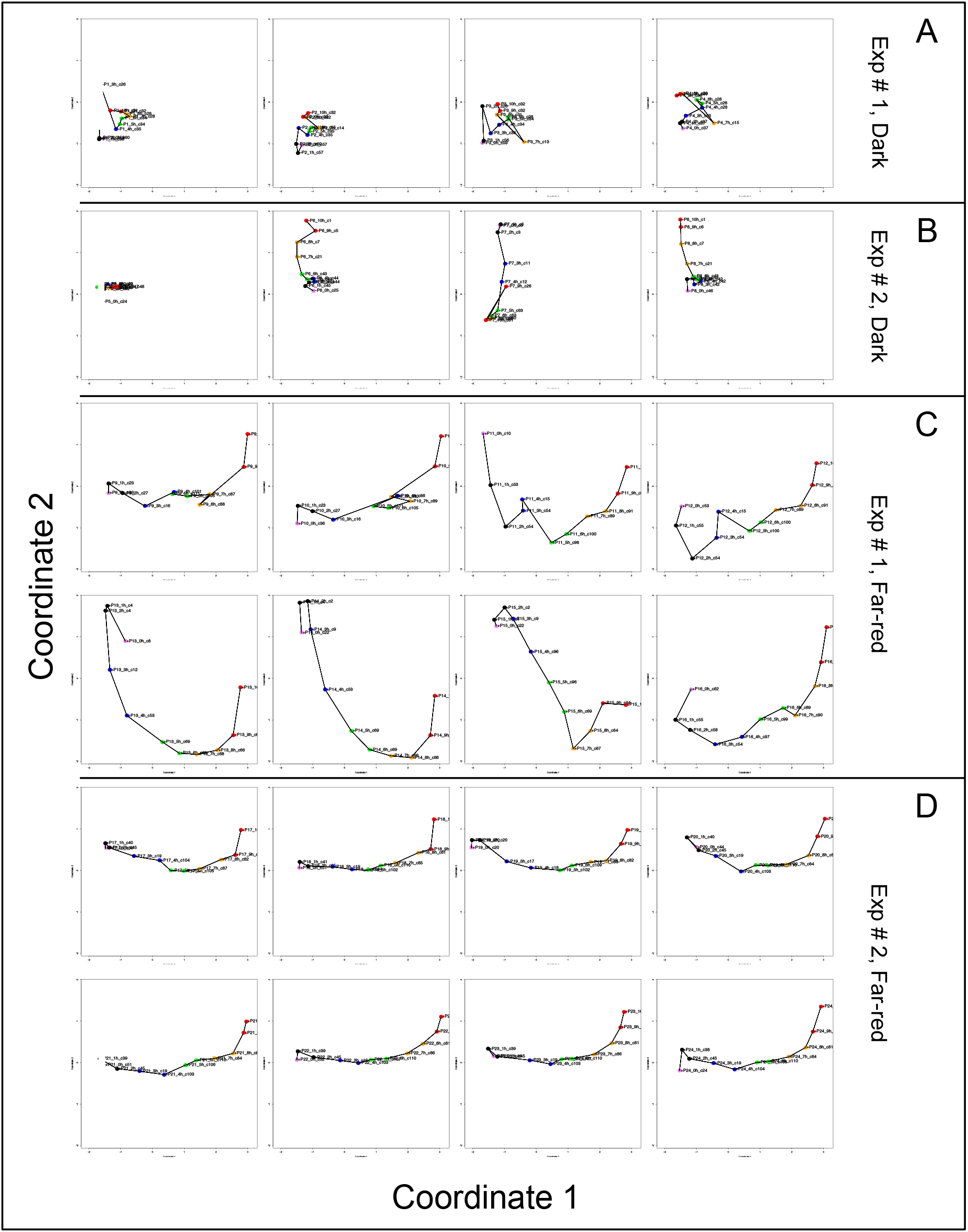
Single cell trajectories of gene expression displayed after multidimensional scaling (MDS) of the expression patterns. Panels (A) and (B) show the trajectories of unstimulated cells of experiment #1 and experiment # 2, respectively. Panels (C) and (D) show the trajectories of far-red stimulated cells of the two experiments. Each data point represents the gene expression pattern of a cell at a given time point at 1h time intervals. The start point (0h) of each trajectory is encoded in pink and the endpoint (10h) in red. The assignment of each data point to a simprof cluster is also given and readable by zooming the pdf version of the figure. The label +P12_1h_c55, for example, indicates that the expression pattern of plasmodium number 12 as measured at 1h after the start of the experiment (corresponding to the onset of the far-red light stimulus in light-stimulated cells) was assigned to Simprof cluster number 55 and that the plasmodium had sporulated (+) in response to the stimulus (+, sporulated; -, not sporulated). All plots are displayed at the same scale.

**Figure 3.**
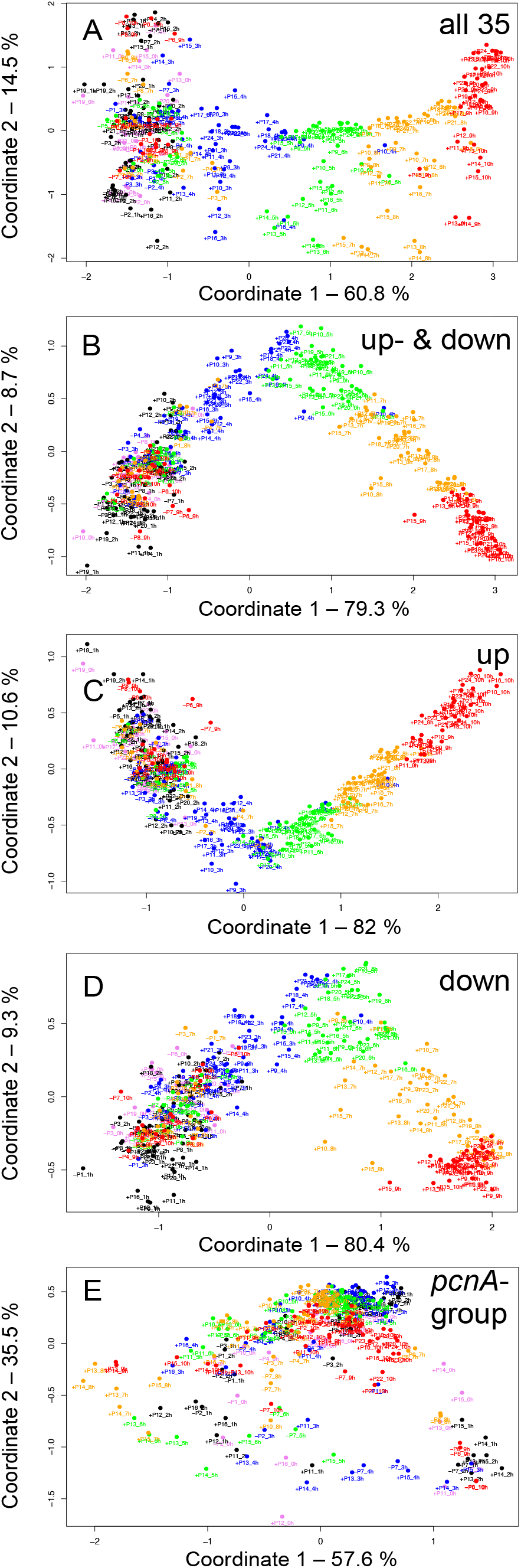
Gene expression patterns of all analysed cells displayed for different sub sets of genes. Multidimensional scaling was performed for the complete set of 35 genes (A), the subset of up- and down-regulated genes (B), or exclusively for up-regulated (C), down-regulated (D) or the *pcnA-group* of genes (E). Each data point represents the expression pattern of an individual cell at a given time point. Time is encoded by color (0h, pink; 1h, 2h, black; 3h, 4h, blue; 5h, 6h, green; 7h, 8h, ocher; 9h, 10h, red). The percent of variance is given for each coordinate.

### Construction and graph properties of Waddington landscape Petri nets

For a further analysis, we performed hierarchical clustering of the expression data for all assayed 35 genes,_differentiation marker and reference genes (SI Table 2), and identified significantly different clusters of expression patterns with the help of the simprof algorithm (Clarke et al. 2008) (SI Fig 5). The temporal sequences of gene expression patterns classified as Simprof significant clusters defined a trajectory for each individual cell and revealed significant differences between cell trajectories (Table 2). To relate gene expression states and trajectories we constructed a Petri net (bipartite graph) as previously described (Rätzel et al. 2020; Werthmann and Marwan 2017), by representing each gene expression state by a *place* and the temporal transit between two states by a *transition* (Fig 4). A single token marking one place of the Petri net indicates the current gene expression state of a cell. The token moves from its place to a downstream place when the transition, connecting the two places through directed arcs, fires. As each transition is connected to exactly two places (one preplace and one post-place), tokens are neither formed nor destroyed when moving through the net, so the gene expression state of the cell remains unequivocally defined at any time. The coherent Petri net obtained this way represents a state machine predicting possible developmental trajectories in terms of Markov chains of gene expression states (Rätzel et al. 2020).

**Table 2.** Single cell trajectories of gene expression. Trajectories are displayed as temporal sequences of gene expression states. Each state is given by the cluster ID number to which it was assigned by the Simprof algorithm. The two experiments, Exp #1 and Exp #2, were performed on two different days, respectively, with the same strain (LU897 × LU898) and under virtually identical experimental conditions.

| Experiment | Cell | 0.0h | 1.0h | 2.0h | 3.0h | 4.0h | 5.0h | 6.0h | 7.0h | 8.0h | 9.0h | 10.0h |
| --- | --- | --- | --- | --- | --- | --- | --- | --- | --- | --- | --- | --- |
| Exp #1, Dark | P1 | 60 | 56 | 59 | 26 | 35 | 34 | 34 | 28 | 28 | 32 | 31 |
| Exp #1, Dark | P2 | 57 | 57 | 60 | 63 | 35 | 35 | 35 | 14 | 32 | 32 | 32 |
| Exp #1, Dark | P3 | 59 | 56 | 26 | 36 | 34 | 34 | 28 | 13 | 31 | 32 | 32 |
| Exp #1, Dark | P4 | 37 | 37 | 37 | 33 | 28 | 28 | 28 | 15 | 29 | 30 | 30 |
| Exp #2, Dark | P5 | 24 | 52 | 52 | 52 | 49 | 52 | 52 | 52 | 48 | 47 | 48 |
| Exp #2, Dark | P6 | 25 | 45 | 45 | 44 | 44 | 43 | 43 | 21 | 7 | 5 | 1 |
| Exp #2, Dark | P7 | 3 | 3 | 3 | 11 | 12 | 63 | 63 | 60 | 60 | 26 | 61 |
| Exp #2, Dark | P8 | 46 | 45 | 45 | 42 | 42 | 42 | 43 | 21 | 7 | 6 | 1 |
| Exp #1, Far-red | P9 | 33 | 23 | 27 | 16 | 101 | 111 | 105 | 87 | 88 | 70 | 79 |
| Exp #1, Far-red | P10 | 36 | 23 | 27 | 16 | 88 | 111 | 105 | 89 | 80 | 75 | 78 |
| Exp #1, Far-red | P11 | 10 | 53 | 54 | 54 | 15 | 98 | 100 | 89 | 91 | 93 | 71 |
| Exp #1, Far-red | P12 | 53 | 55 | 54 | 54 | 15 | 100 | 100 | 89 | 91 | 92 | 71 |
| Exp #1, Far-red | P13 | 8 | 4 | 4 | 12 | 53 | 69 | 69 | 68 | 66 | 65 | 95 |
| Exp #1, Far-red | P14 | 22 | 4 | 2 | 9 | 53 | 69 | 69 | 68 | 66 | 65 | 95 |
| Exp #1, Far-red | P15 | 22 | 4 | 2 | 9 | 96 | 96 | 69 | 67 | 64 | 94 | 95 |
| Exp #1, Far-red | P16 | 62 | 55 | 58 | 54 | 97 | 99 | 89 | 90 | 93 | 75 | 78 |
| Exp #2, Far-red | P17 | 50 | 40 | 45 | 19 | 104 | 103 | 105 | 87 | 82 | 72 | 79 |
| Exp #2, Far-red | P18 | 51 | 41 | 25 | 19 | 104 | 102 | 110 | 85 | 81 | 72 | 77 |
| Exp #2, Far-red | P19 | 20 | 20 | 20 | 17 | 18 | 102 | 109 | 83 | 82 | 72 | 77 |
| Exp #2, Far-red | P20 | 44 | 40 | 45 | 19 | 103 | 108 | 110 | 84 | 82 | 73 | 77 |
| Exp #2, Far-red | P21 | 51 | 39 | 45 | 19 | 103 | 106 | 110 | 84 | 82 | 74 | 76 |
| Exp #2, Far-red | P22 | 52 | 39 | 45 | 19 | 103 | 107 | 110 | 86 | 81 | 73 | 76 |
| Exp #2, Far-red | P23 | 46 | 39 | 45 | 19 | 103 | 107 | 110 | 86 | 81 | 73 | 77 |
| Exp #2, Far-red | P24 | 24 | 38 | 45 | 19 | 104 | 108 | 110 | 84 | 81 | 74 | 77 |

**Figure 4.**
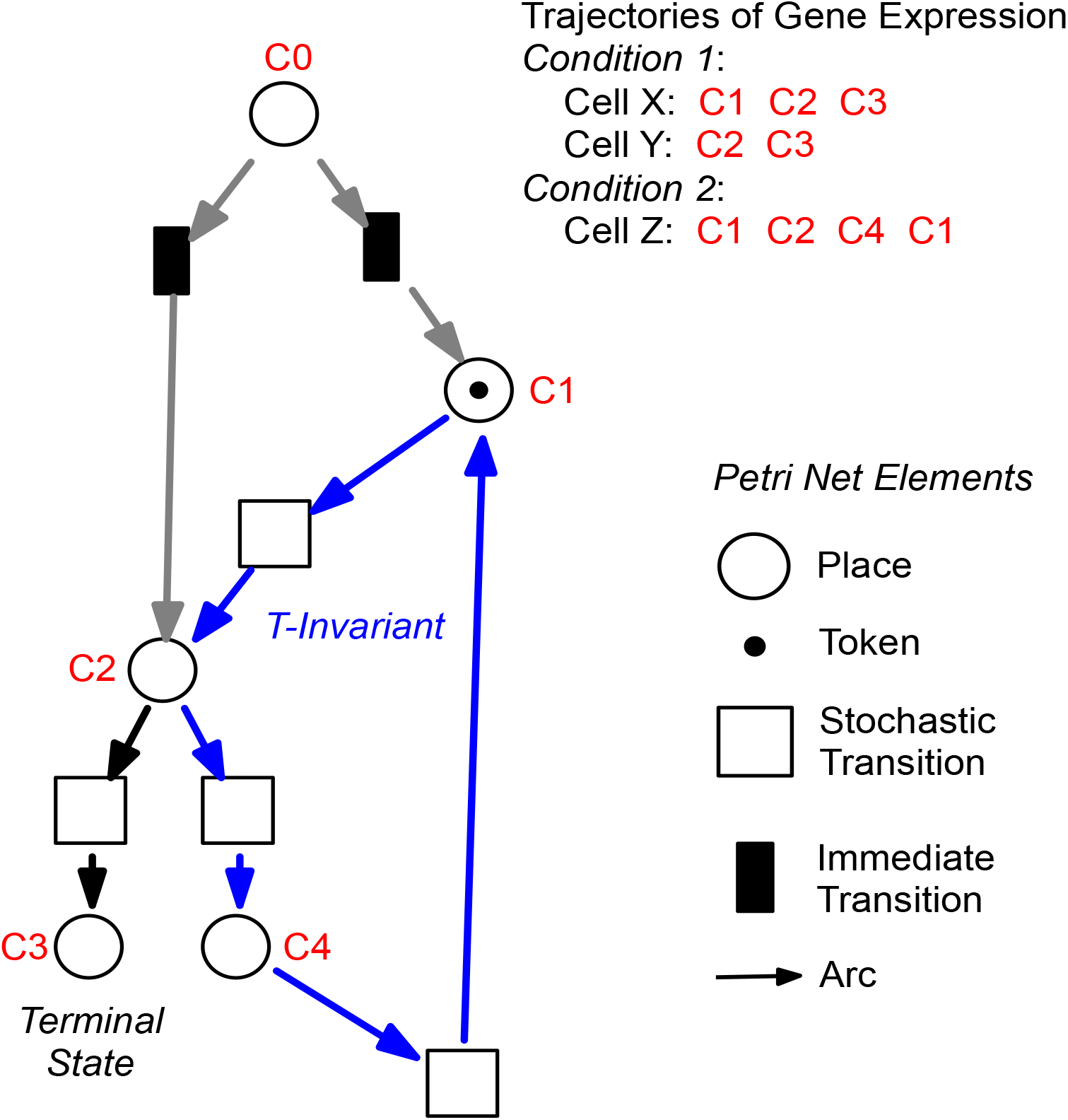
Construction of Petri nets from single cell trajectories of gene expression. Petri nets are directed, bipartite graphs with two types of nodes, places and transitions, that are connected by arcs (see symbols, lower right). Petri nets are used in this work to model state machines, as exemplified in the following. Any gene expression state of a cell, as defined by its assignment to a Simprof significant cluster of gene expression patterns, is represented by a corresponding place (drawn as a circle). Any transit between two states is mediated by a transition (drawn as a rectangle). The current gene expression state of a cell is indicated by one token which marks the respective place. When a place contains a token, one of its posttransitions can fire to move the token into its post-place. (A post-transition of a place is a downstream transition which is immediately connected to that place, as indicated by a directed arc). Because each transition of the Petri nets as they are used here, has exactly one pre-place (one incoming arc) and one postplace (one outgoing arc), and because all arc weights are one, tokens can neither be produced nor destroyed, and the state of gene expression remains unequivocally defined. A transit cycle, *i.e*. the ensemble of reactions that bring a subsystem back to the state from which it started, is called transition-invariant (T-invariant) (Sackmann et al. 2006). The arcs that contribute to the T-invariant of the Petri net displayed in the figure are highlighted in blue. Petri nets, as they are used in this paper, contain one additional place C0, which does not represent a gene expression state. For simulation, C0 defines the initial gene expression state of the cell by randomly delivering its token to one of the places that are connected to C0 through so-called *immediate transitions* (filled in black) that fire immediately when the simulation starts (for details see (Rätzel et al. 2020)). Connection to C0 also graphically highlights the places representing those gene expression states in which cell trajectories started. In the example shown, the cell trajectory started in a gene expression state assigned to Simprof cluster C1. The token can move to C2 where it randomly moves to either C3 (a terminal state in this example) or to C4, from which it may return to C1 and possibly continue.

The basic modelling principles are summarized in Table 3. We observe the following structural properties of the model which we call ‘*Waddington landscape Petri net*’:

- Each transition has exactly one pre-place and one post-place.
- There are places having more than one post-transition. These post-transitions are in conflict. But, because every transition has exactly one preplace, each conflict is a free choice conflict, meaning the token is free to choose which route to take, predicting a corresponding free choice for the cell (see Discussion).
- There are places having more than one pre-transition, i.e. alternative paths may rejoin. Thus, the Petri net structure does not form a tree.
- There are cycles: a cell may switch back to previous states or oscillate between states as defined by the expression patterns of the set of observed genes.

**Table 3.**
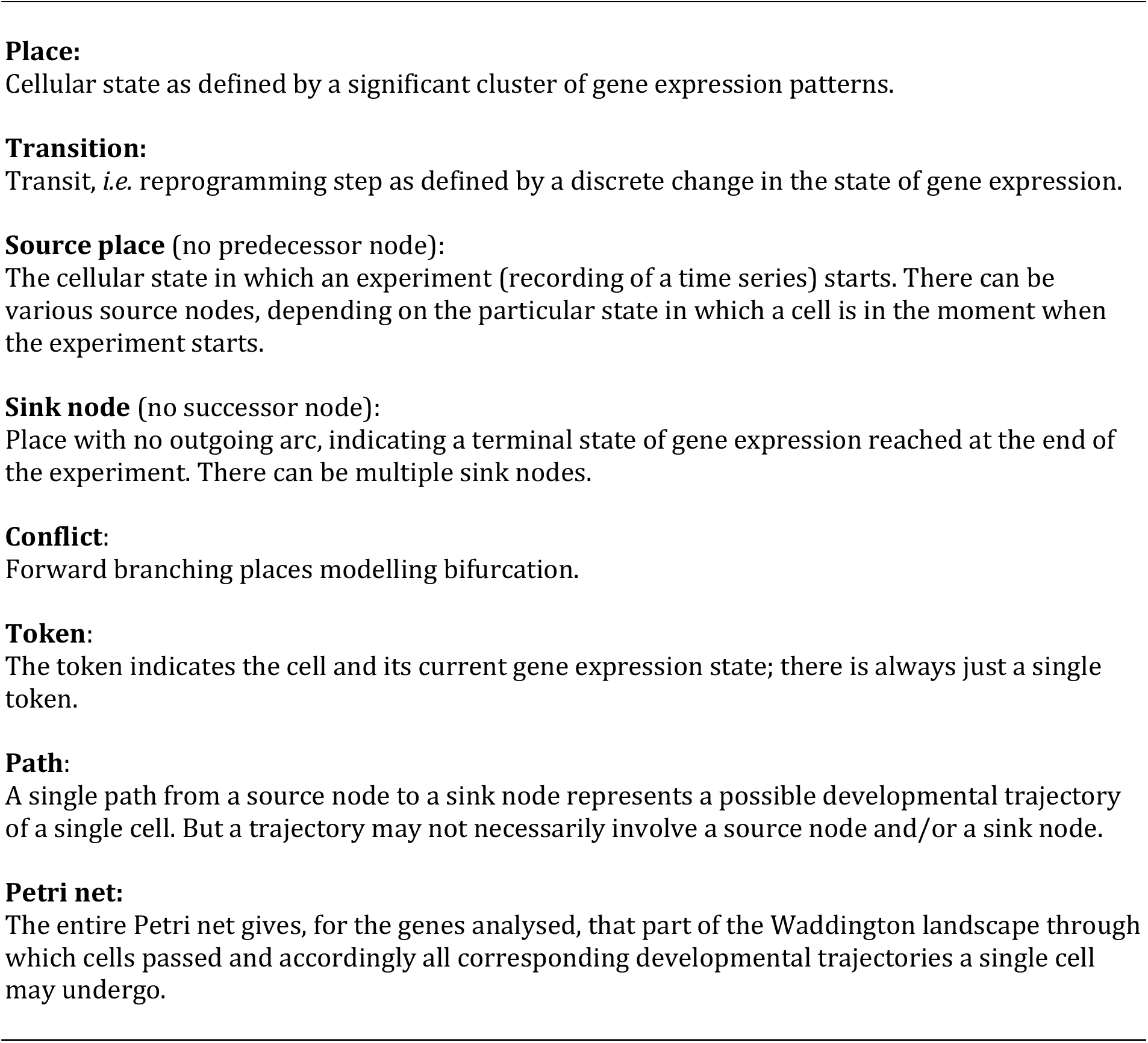
Terminology, basic modelling principles, and Petri net elements

For technical reasons we add immediate transitions starting alternative trajectories, in order to get a statistical distribution of states in which the experiments have started or will start with a given probability.

In contrast to most state-of-the-art pseudo-time series approaches found in the literature (Saelens et al. 2019), the structure of the Waddington landscape Petri net is not restricted to a partial order, meaning it is neither restricted to a directed acyclic graph nor to a tree. Instead we obtain what is known in Petri net theory as ‘*state machine*’, also called in other communities ‘finite state machine’ or ‘finite automata’, which may involve cycles.

A state machine with one token and its reachability graph, or Markov chain for stochastic Petri nets, are isomorph (*i.e*. have the same structure, there is a 1-to-1 correspondence); to put it differently: our (stochastic) Petri net represents the Markov chain of states the cells assume in the course of their developmental trajectory and accordingly on their walk through the Waddington landscape. We assume that the Petri net represents the corresponding region of the Waddington landscape predicting possible developmental paths a single cell can follow, which of course yields a state machine.

Representing Markov chains as Petri nets comes with a couple of advantages. First, Petri nets are equipped with the concept of T-invariants, which belong to the standard body of Petri net theory from very early on (Lautenbach 1977). We consider T-invariants as crucial in terms of biological interpretation of the generated net structures (Heiner 2009; Sackmann et al. 2006). The computation of T-invariants is rather straightforward for state machines; due to their simple structure it holds:

- each cycle in a state machine defines a T-invariant, and
- each elementary cycle (no repetition of transitions) is a minimal T-invariant.

Second, modelling the differentiation-inducing stimuli, what we have not done so far, would turn some of the free choice conflicts into non-free choice conflicts, which involves, technically speaking, leaving the state machine net class. To unequivocally identify transits that are stimulus-dependent, we need a higher data density which we will hopefully achieve in one of our next experiments. With stimulus-dependent transitions, the constructed Petri nets and their Markov chains do not coincide anymore, instead the Markov chains as well as the reachability graph are directly derived from the Petri nets and may be analyzed by standard algorithms. Finally, our Petri net approach paves the way for the actual ultimate goal of our future work – reconstructing the underlying gene regulatory networks based on the reachability graphs encoded by the Waddington landscape Petri nets.

### Characterization of Petri nets constructed from gene expression data

Figure 5 displays a Petri net assembled from cell trajectories considering the set of 35 genes. The graphical representation laid out using the Sugiyama algorithm emphazises the directionality of concurrent processes (Sugiyama et al. 1981). The net indicates that cells started in different states (connected to C0; see Legend to Fig 4 for details) and, after stimulation by far-red light, proceeded to a small set of terminal states (places C71, C76, C77, C78, C79, C95) via multiple, more or less highly connected intermediate states. To facilitate the interpretation of the Petri net, we have colored the places and transitions according to different criteria. In Fig 5A, the transitions being specific to cells of experiments #1 or #2 and for dark controls or light-stimulated cells in the respective experiments, are colored differently. Each place is colored according to the relative frequency of its corresponding gene expression state, indicating that some states occurred more frequently than others. From this representation it is obvious that cells of experiments #1 and #2 form different branches, in part projecting onto different terminal states, reflecting accordingly different developmental trajectories to commitment and sporulation. Figure 5B displays the same Petri net, but with a different color coding. Here, transitions are colored according to how frequent the corresponding transits occurred, indicating that some paths were more frequently taken than others. Places are colored according to their relative stability, defined as the average residence time of a cell in the respective state (see Methods for details). Coloring indicates that cells reached a terminal state through states of different stability, e.g. meta-stable intermediates.

**Figure 5.**
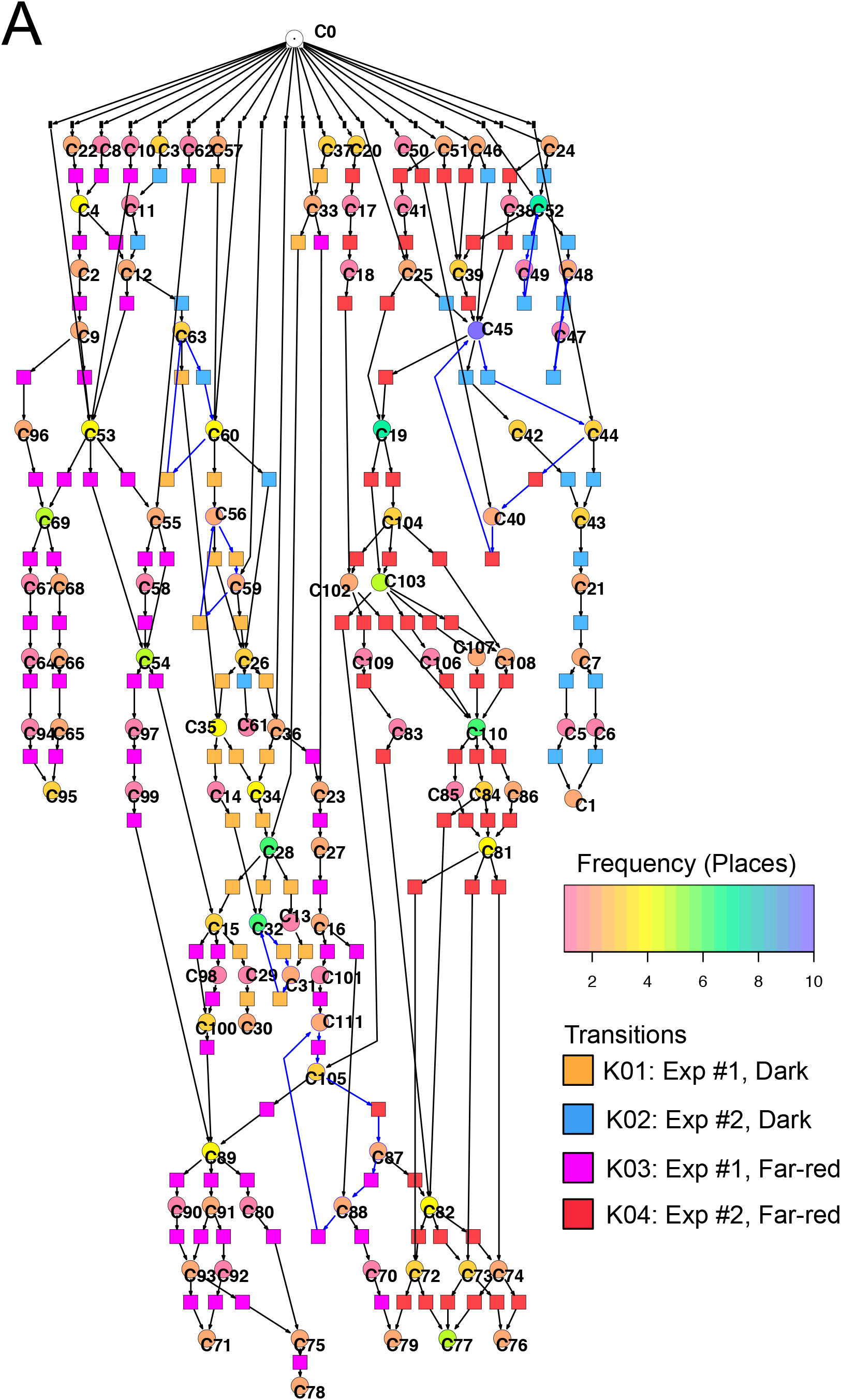
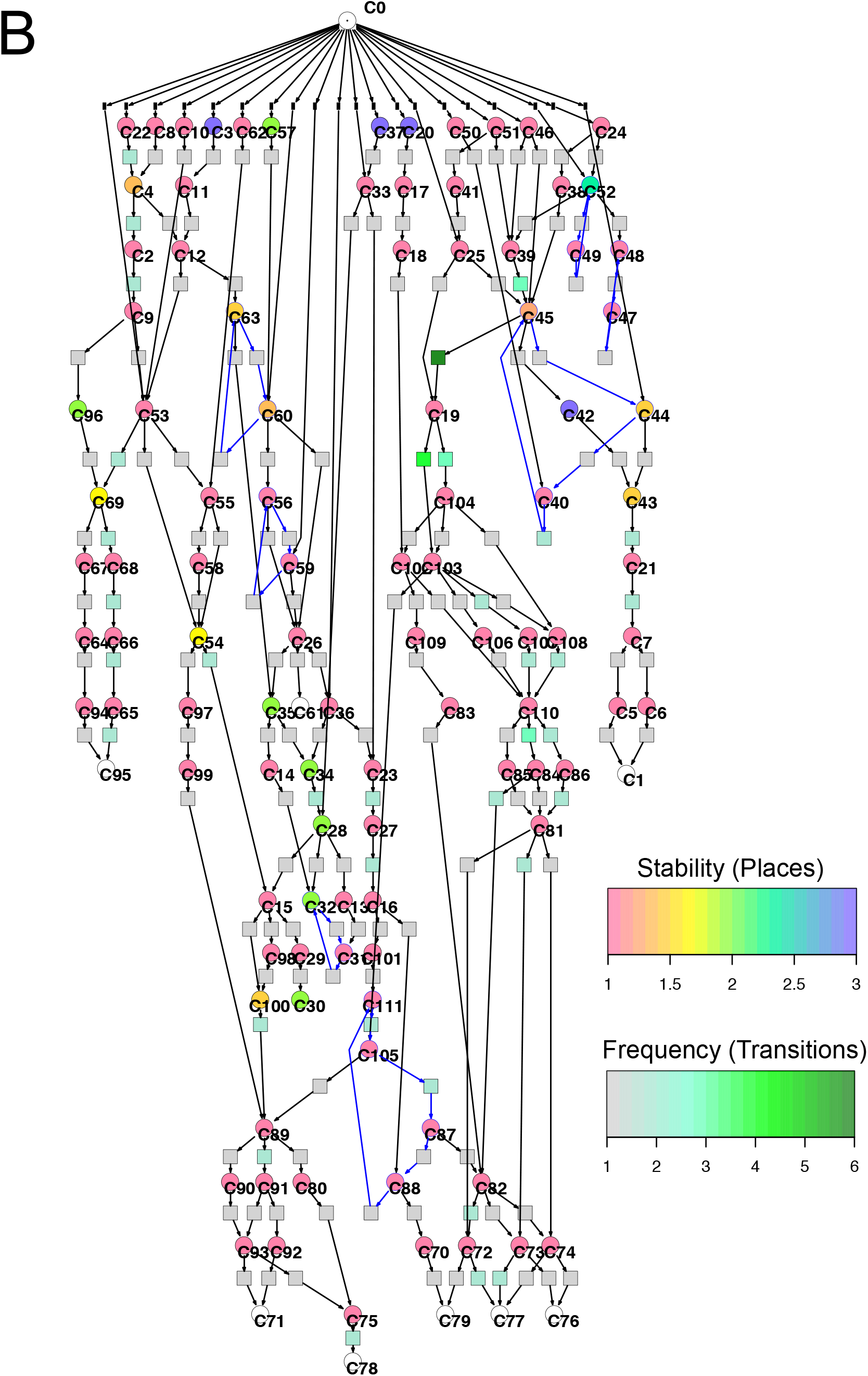
Two copies of the same Petri net, automatically constructed from the single cell trajectories of gene expression as displayed in Table 2. Places and transitions in the two nets were colored according to different criteria. Arcs as part of T-invariants are highlighted in blue. (A) As indicated by the rainbow color key, places are colored according to the relative frequency with which respective gene expression states occurred in the data set. Transitions are colored according to whether corresponding transits occurred in experiments #1 or #2, and whether cells were far-red stimulated or un-stimulated (dark controls), respectively, as indicated by the panel lower right. (B) Places are color coded according to the relative stability of the states of gene expression they represent. This relative stability indicates how long a cell on average resided in a certain state. The color of transitions indicates, in absolute numbers, how frequent a corresponding transit occurred in the data set.

Un-stimulated cells (Table 2, dark controls) spontaneously switched between significantly different states of gene expression. Their trajectories gave three disconnected Petri nets (Fig 6A; the three nets were connected to C0 for technical reasons, see legend to Fig. 4).

**Figure 6.**
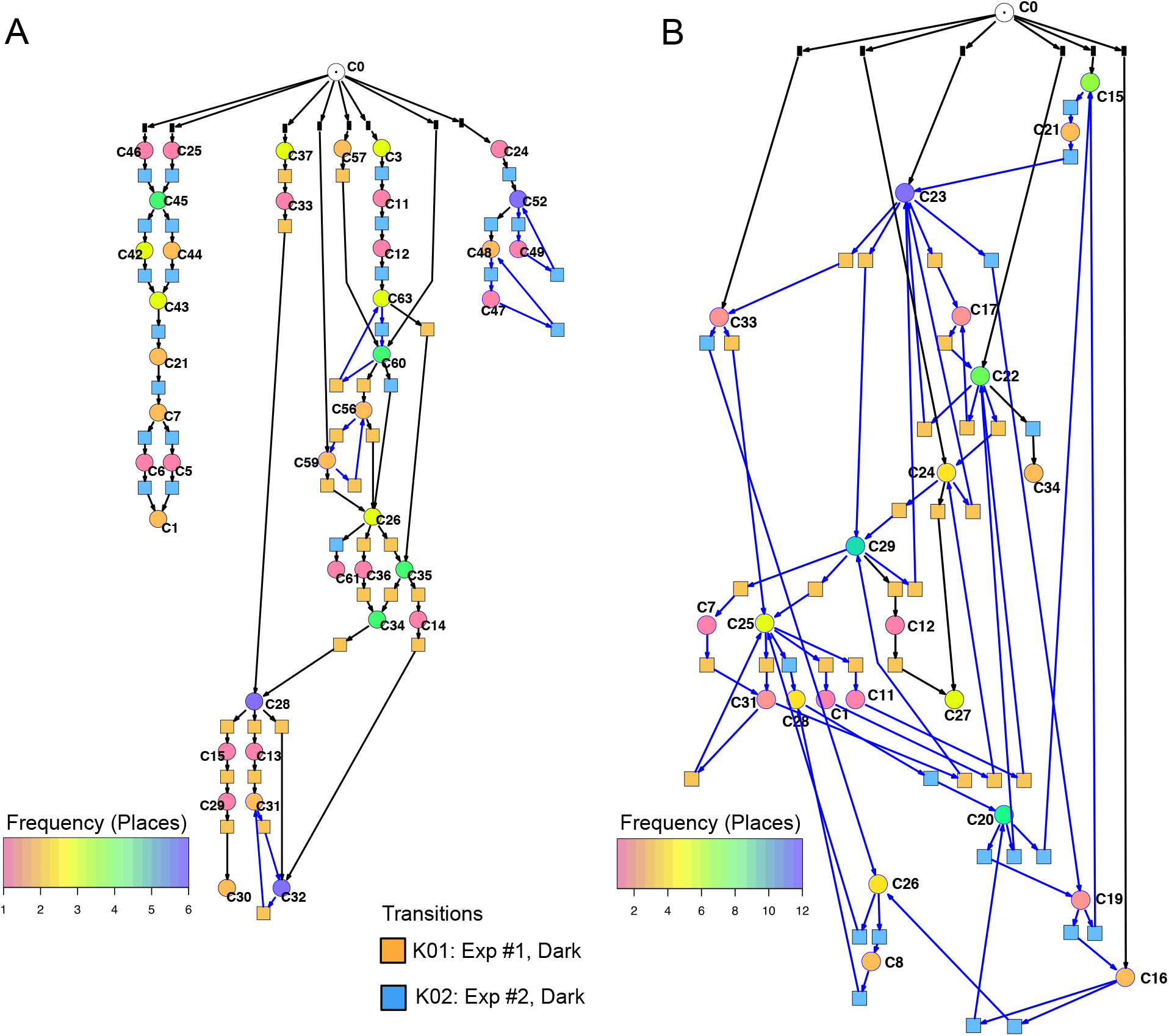
Petri nets constructed for different sets of genes from the gene expression trajectories of un-stimulated cells. (A) Trajectories considering the complete set of 35 genes yield three disconnected Petri nets containing a low number of T-invariants (as indicated by arcs highlighted in blue). (B) When only those genes are considered that are up- or down-regulated in response to far-red stimulation (omitting the *pcnA*-group of genes), the trajectories of un-stimulated cells give one coherent Petri net which is almost covered with T-invariants. Color coding of places indicates the relative frequencies of states of gene expression in un-stimulated cells. Differences in gene expression patterns of un-stimulated cells between the cells of experiment #1 and #2 are indicated by the appearance of experiment-specific transits and accordingly differently colored transitions.

Petri nets of Fig 5A,B and Fig 6A indicate that the expression pattern in both, stimulated and un-stimulated cells developed predominantly in forward directions while there were some transits back to previous states creating so-called transition-invariants (T-Invariants; Fig 4). We asked whether temporal expression differences in the *pcnA-group* of genes might have added to this directedness. Therefore, we constructed a Petri net from trajectories based on significant clusters, this time exclusively clustering the subset of up- and down-regulated genes. Basic features found in the Petri nets of up- and down-regulated genes were similar to the ones found in the Petri nets for the full set of 35 genes: Trajectories formed parallel main branches, there were intermediate nodes of different stability, of different connectedness, and hence states that occurred with different frequency (SI Fig 6). In contrast, there was a high number of minimal T-invariants that heavily involved places representing gene expression states that occurred in un-stimulated cells. A Petri net built by considering only transits that occurred in un-stimulated cells (Fig. 6B) was nearly covered with T-Invariants, indicating spontaneous, reversible alterations in the expression of up- and/or down-regulated genes. Again, states of gene expression displayed different stability. Considerable variation in the expression level of the up-and down-regulated genes is even most obvious from the heat map of initial states from which trajectories emerged and of terminal states that were observed during the experiment (Fig 7). The high density of T-invariants in the un-stimulated cells suggests that similarly, the T-invariants involving places corresponding to light-stimulated cells are due to gene expression changes that do spontaneously occur before cells are caught by a new attractor formed in response to the far-red stimulus (see Discussion).

**Figure 7.**
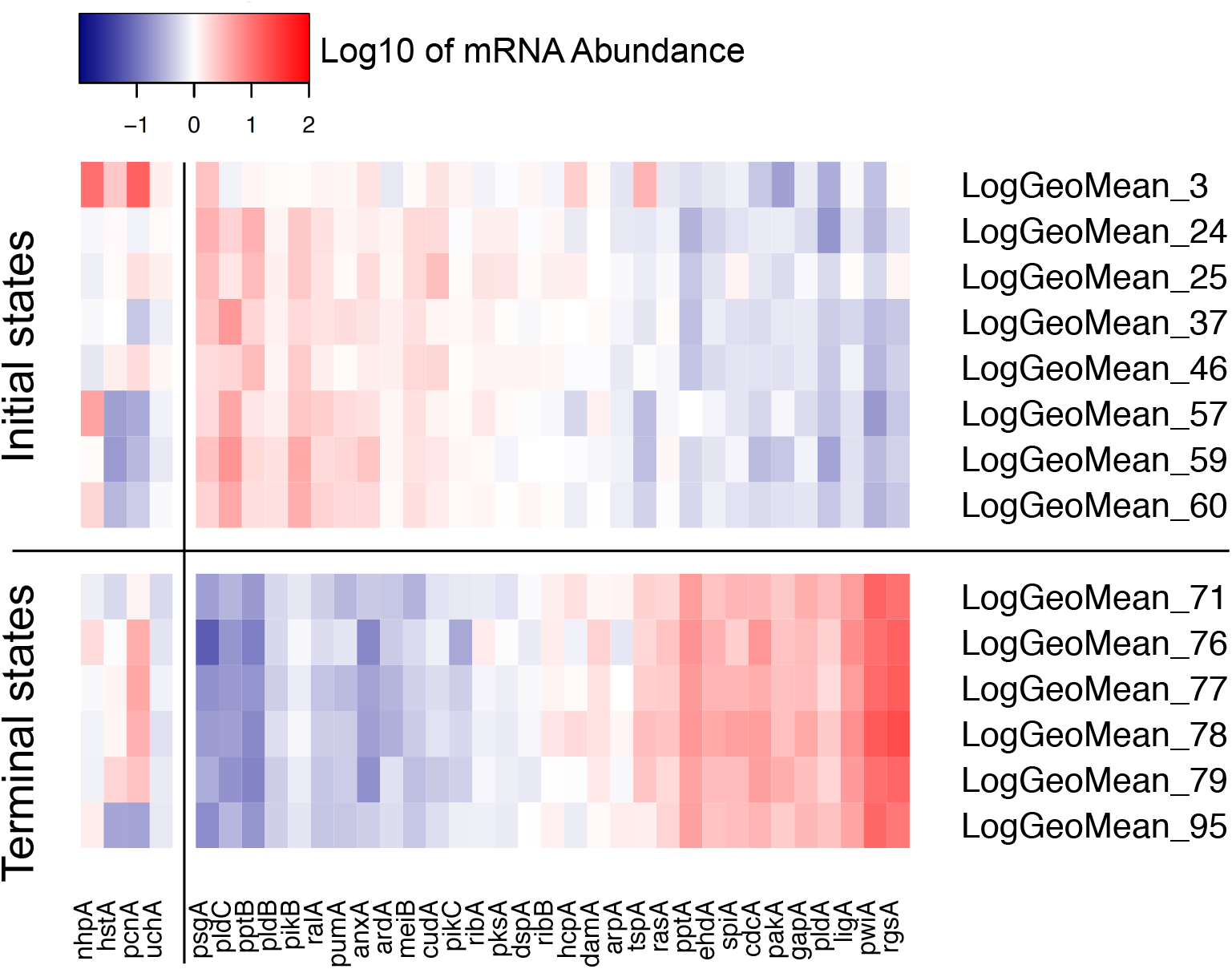
Heat map visualizing the variability of initial and terminal states of gene expression. The initial states displayed in this panel are the start points of trajectories of un-stimulated cells as listed in Table 2. The terminal states are endpoints of trajectories of far-red stimulated cells corresponding to terminal places of the Petri net of Fig. 5. Color-coded expression values correspond to the logarithm to the base 10 of the geometric mean of all expression values of a respective cluster with its ID number indicated on the right side of the panel. For clarity, expression of the *pcnA-group* of genes is displayed as a separate block.

Fig 6 also shows that selecting sets or subsets of genes for hierarchical clustering and subsequent Petri net construction may yield Petri nets of different structure delivering accordingly non-redundant information on corresponding subsets. This is also shown in Table 4 for the set of 35 genes and for subsets, the down-regulated, up-regulated, down- and up-regulated, and the *pcnA*-group of genes. Except for the *pcnA*-group, the number of places per gene was approximately the same. The number of transitions per gene however became less with more genes considered. This suggests that up-regulation, down-regulation and even expression of the *pcnA*-group of genes are at least partly coordinated or co-regulated processes. The average number of minimal T-invariants per gene compared for the different groups of genes (Table 4) suggests that reversibility observed for the subsets vanishes when more genes are considered, obviously due to combinatorial effects.

**Table 4.**
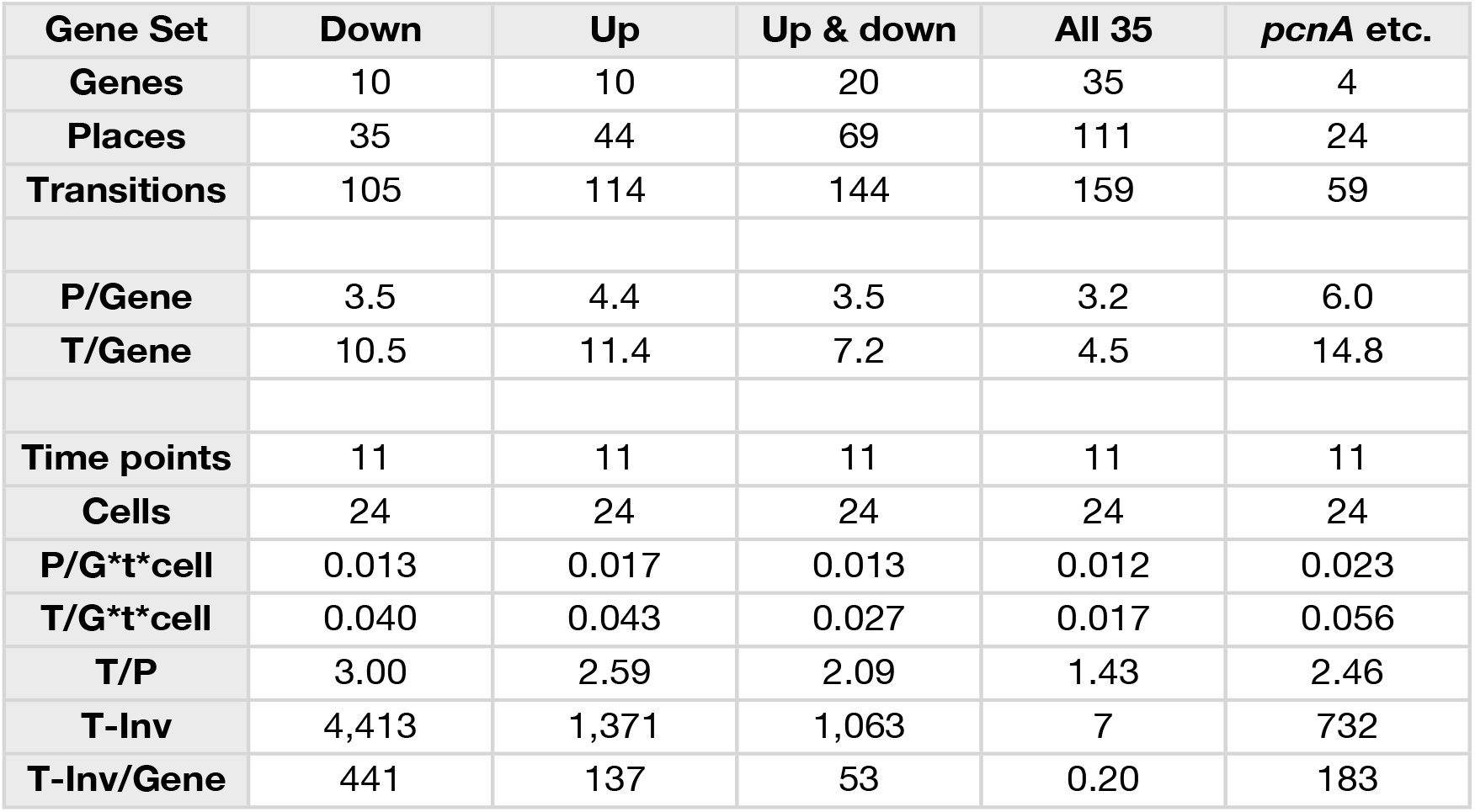
Number and relative frequency of places, transitions, and T-invariants of Petri nets constructed for different subsets of genes (see legend to SI Figure 4) from the data of cells listed in Table 2.

### Single cell trajectories reveal qualitatively different patterns in differential gene regulation

We have argued that Petri net modelling disentangles the complex response of cell reprogramming (Rätzel et al. 2020) and predicts feasible developmental pathways through the Waddington landscape, resulting in significantly distinct single cell trajectories. To reveal similarities and differences of the expression kinetics of individual genes, we plot the geometric mean of the concentration values of the mRNA in a cluster logarithmically, normalized to its concentration at the start of the experiment (*t*=0h) as a function of time for any single cell trajectory. In this kind of plot, the time course of the mRNA of each gene starts at the same point, while the slope of the curve indicates the x-fold change in mRNA abundance over time. Plotting subsets of genes suggests that trajectories through different regions of the Petri net of Fig 5 indeed emerge from qualitatively different expression kinetics, and that genes are also differently regulated relative to each other when different trajectories are compared. In the example shown in Fig. 8, *pldA* is early up-regulated in quite a number of trajectories, followed by *pwiA* and finally by *ligA* and *rgsA* that appear strongly correlated at least in some of the plots. Qualitatively different patterns of regulation relative to each other are also evident for the three phospholipase D-encoding genes (SI Fig 7). The *pldA* gene is up-regulated while *pldB* and *pldC* are down-regulated. In some of the trajectories, the initial change in the concentration of the *pldA* and *pldC* mRNAs is inverse as compared to the overall time course. A more comprehensive representation with more genes displayed makes similarities and differences between trajectories even more obvious (SI Fig 8). Here, we observe a phenomenon, which is also seen in Table 2, namely that cells remain in a certain state for some time. This occurs predominantly in the unstimulated cells but is also seen in some of the light stimulated cells, e.g. in those that proceed to state C95 (Fig 5A,B). Cells seemingly are trapped in a meta-stable state (e.g. C96, C69, C100 etc.; Fig 5B) for some time until the developmental program proceeds. We presumably will need more data to see whether this is an artefact which occurs by the discretization of gene expression through clustering. Conversely, discretization might help to identify tipping points for the differential regulation of gene expression as the plots in SI Fig 8 suggest.

**Figure 8.**
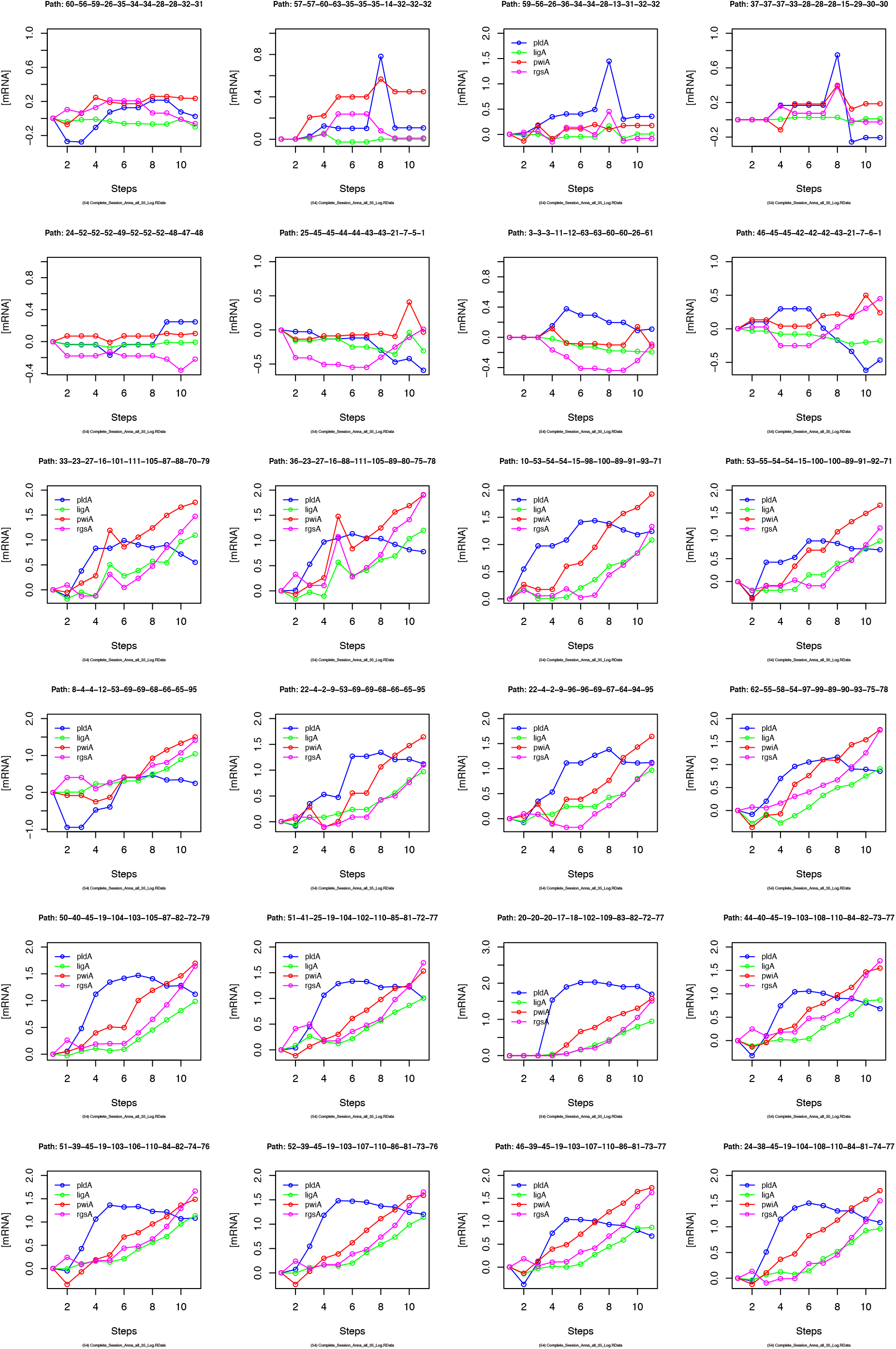
Gene expression kinetics as derived from single cell trajectories. Each panel represents the trajectory of one single cell as characterized by subsequent states of expression of the same, arbitrarily chosen set of genes. For each gene, the logarithm to the base 10 of the geometric mean of the expression values of each cluster was plotted against time. Plotting the geometric mean instead of individual expression values results in discretization of the data while comparing different trajectories.

## Discussion

We have analysed the gene expression dynamics in response to a differentiation-inducing stimulus pulse in true time by repeatedly taking samples of large, multinucleate plasmodial cells. Control experiments have demonstrated that the gene expression patterns in samples simultaneously retrieved from different sites of a large plasmodial cell did not deviate within the range of the technical accuracy of the measurements. This again confirms that the plasmodial cytoplasm, at least at the level of macroscopic sampling, can be considered as a homogeneous reaction volume. Accordingly, repeated sampling of the same plasmodium allows for the analysis of true time series of gene expression at the level of an individual multinucleate cell. The cytoplasm containing millions of nuclei is continuously mixed by the vigorous shuttle streaming. Therefore, the observed changes in the state of gene expression over time most likely reflect the (non-linear) dynamics of the system rather than stochastic gene expression noise.

Gene expression states of the cells were defined by hierarchical clustering and discretized by assigning each gene expression pattern to a simprof significant cluster (Clarke et al. 2008; Rätzel et al. 2020). Trajectories of subsequent discrete states were then assembled into a state machine implemented as a Petri net. In the Petri net, each gene expression state is represented by a place and each transit between two states is represented by a transition.

The Petri net, as it has been defined in this and previous studies (Clarke et al. 2008; Rätzel et al. 2020; Werthmann and Marwan 2017), models gene expression trajectories as Markov chains (Gagniuc 2017), which assumes that each subsequent state only depends on the current state of a cell and not on its previous states, *i.e*. it does not depend on the individual history of a cell. This assumption is commonly made by computing pseudotime series from snapshots of individual mammalian cells (Bendall et al. 2014; Chen et al. 2019; Haghverdi et al. 2016; Marr et al. 2016; Setty et al. 2019; Shin et al. 2015; Street et al. 2018; Weinreb et al. 2018). It follows the principle of parsimony in making not more assumptions than necessary and giving the simplest possible explanation for an observed phenomenon. Practically this means that any path which a token can take through the Petri net, by stochastic firing of the transitions, translates into a feasible trajectory of an individual cell. Hence, firing of a transition does only depend on the marking of the preplace of this transition and not on the identity of any upstream places from which the token originally came. Defining the cell’s state of gene expression by measuring more genes might well diversify places and hence change the structure of the Petri net. This has been demonstrated by constructing nets from subsets of genes. In the examples provided, the structure of the Petri net changed and the number of T-invariants increased drastically upon reduction of the number of considered genes (Table 4).

If the structure of the Petri net does depend on the set of genes analysed, what is its actual value? The actual value is that it reveals the behavior of states defined by sets or subsets of genes. Limiting the analysis to the chosen subset of up- and down-regulated genes, as we have done here, revealed extensive on- and off-switching of the genes in unstimulated cells that are differentially regulated in response to a differentiation-inducing stimulus. This became immediately obvious through structural analysis of the Petri net by determining the number of minimal T-invariants. Displaying the net in Sugiyama representation revealed another phenomenon with respect to this subset of genes. The light stimulus caused directed development towards a small number of terminal states reducing the overall number of alternative states in which the cells resided. This suggests that a cellular attractor is formed in response to the stimulus causing the commitment to differentiation.

Coloring the transitions of the Petri net according to the frequency by which transits occurred, allows identification and visualization of main paths, *i.e*. paths which the system preferably took. Coloring places according to the relative stability of the states they represent indicated metastable states that were not necessarily identical to highly connected places. Places having many pre-transitions (many incoming arcs) represent states, the system is likely to assume, like a corrie in the metaphor of the Waddington landscape, through which the system will pass. Places having many post-transitions (many out-going arcs) represent branching points from which the system has multiple options to proceed.

Our analysis has confirmed former observations (Rätzel et al. 2020), now at considerably larger resolution in time, that unstimulated cells spontaneously and reversibly change their expression pattern. These changes involved the expression of genes that are differentially regulated in response to a differentiation-inducing stimulus. Spontaneous switching of gene expression patterns is at least one reason why stimulated cells started their way to commitment and differentiation from quite different states, indicating substantial heterogeneity in the population of cells. In other words, cells can start differentiation while being in various different states. The differentiation-inducing stimulus then collects or focusses these cells onto a narrow set of states like an attractor of a dynamic system would do. This phenomenon is graphically revealed by the funnel- or cone-like appearance of the Petri net in the Sugiyama layout (SI Fig 6).

The response of a cell to a differentiation-inducing stimulus seems to depend on the cell’s current internal state. Fig 5A revealed distinct main branches (visible through transitions of different color) for cells from experiment #1 as compared to experiment #2, suggesting that the response of the cell in terms of its developmental pathway did indeed depend on the initial physiological or gene expression state in which the cell resided while receiving the stimulus. The cells proceeded to slightly different terminal states that however might belong to the same cellular attractor.

One might be tempted to suspect a certain structure in the list of subsequently recorded trajectories (Table 2). Changes in the intial state of the plasmodia until the time of stimulus application might have occurred as the experiment proceeded. Similarities in subsequently recorded trajectories may be by chance and we cannot draw any final conclusion because the number of analysed plasmodia is by far too low. With more plasmodia analysed and more genes measured, we might discover that the individual history of a cell indeed matters, meaning that the Markov assumption is wrong. Even if this should be the case, the Petri net representation would still be valid, however with the firing probability of certain transitions depending on which path the token came from. Technically, this dependency could be implemented in the form of a colored Petri net. In this context it is trivial and at the same time important to note that the identity of a place (i.e. state) is always defined by the set of genes that have been measured. Measuring more genes might split any place into more places or even into a separate Petri net with an increased overall number of places. This holds not only for states defined by gene expression but also for cellular states defined by the covalent modification of proteins *etc*.

We have previously argued that the Petri net depicts aspects of the topology of the Waddington landscape (Huang et al. 2009; Waddington 1957) with respect to and limited to the set of observed genes (or measured molecular entities) (Rätzel et al. 2020; Werthmann and Marwan 2017). Then, the token in the Petri net corresponds to the marble rolling down the Waddington landscape as developmental processes unfold. Each Petri net place represents, albeit implicitly, a significantly distinct gene expression state. Despite this implicit representation, the temporal information on each gene for each cell trajectory is available and we have used this information to reveal the temporal hierarchy of differentially regulated genes. The Petri net representation disentangles accordingly the complex gene expression response and identifies alternative regulatory programs or routes. Using this information, the underlying regulatory network can be inferred by applying appropriate algorithms (Durzinsky et al. 2013; Durzinsky et al. 2011; Marwan et al. 2008). This is a possible next step to go.

## Materials and Methods

### Plasmodial strain, growth of cells, sample preparation, and gene expression analysis

Sporulation-competent plasmodial cells of wild type strain LU897 × LU898 (Starostzik and Marwan 1998) were obtained as previously described (Rätzel et al. 2020; Starostzik and Marwan 1998). A total of 2.8 gram of plasmodial mass was applied to a 14 cm Ø Petri dish that contained 90 ml of 0.5 × Golderer agar (Golderer et al. 2001), a salt solution of pH 4.6 supplemented with 10 g/L peptone from meat (Sigma Aldrich), 1.5 g/L yeast extract (Becton, Dickinson & Co.), and 1.55 g/L glucose, as well as 0.01% (w/v) niacin, 0.01% (w/v) niacinamide, 0.1% (w/v) CaCO_3_, and 0.14 mM CuCl_2_. After starvation for seven days at 22 °C in complete darkness, sporulation was induced with a 15 min pulse of far-red light (2 ≥ 700 nm, 13 W/m^2^) (Starostzik and Marwan 1998). Before and at one hour time intervals after the start of the far-red pulse, samples were taken in duplicate at randomly chosen, distant positions on the plate. Each sample was obtained by picking an agar plug of 1.13 cm^2^ with the cutted bulb of a disposable Pasteur pipette (EA62.1; Carl Roth, Karlsruhe, Germany). The plasmodial mass on the agar plug was scraped off with a polystyrene spatulum and transferred into a vial of glass beads immersed in liquid nitrogen. After extraction of RNA and removal of contaminating DNA (Marquardt et al. 2017), the relative abundance of the mRNAs of 35 genes, differentiation marker and reference genes (Hoffmann et al. 2012) (SI Table 2) was analysed by gene expression profiling (GeXP), a multiplex RT-PCR method (Hayashi et al. 2007) as previously described (Marquardt et al. 2017; Rätzel and Marwan 2015).

### Data analysis pipeline and automated generation of Petri nets

To correct for differences in the concentration of total RNA and in the efficiency of the RT-PCR reaction, the gene expression values were normalized to the median of the estimated concentrations of mRNAs of the 35 genes in each RNA sample. Each normalized expression value was subsequently normalized to the geometric mean of all values obtained for a given gene, and this was performed separately for each gene.

Data were analysed and processed with a revised and extended pipeline written in R (R Core Team 2016), based on the previously described script (Rätzel et al. 2020). The normalized gene expression data were clustered and significant clusters were determined with the help of the Simprof algorithm (Clarke et al. 2008) as provided by the clustsig package (Whitaker and Christman 2014). Expression patterns were visualized in the form of a heatmap generated by the heatmap.2 function, provided as part of the gplots package (Warnes et al. 2016). Changes in gene expression over time were visualized by multidimensional scaling based on euclidean distance (Gower 1966) with the help of the *cmdscale* function provided as part of the stats package v3.5.1 (R Core Team 2016). Petri nets were constructed from single cell trajectories of gene expression as previously described (Rätzel et al. 2020). Each trajectory is a temporal sequence of gene expression states, where each state corresponds to a Simprof significant cluster. Petri nets specified in ANDL format (Abstract Net Description Language) (Heiner et al. 2013) were imported into Snoopy (Rohr et al. 2010) and graphically displayed by running the Sugiyama layout algorithm (Sugiyama et al. 1981). Petri net places, each representing a gene expression state, were colored according to the relative temporal stability of the expression state or according to the relative frequency with which each gene expression state occurred. Petri net transitions, corresponding to transits between states were colored according to the frequency with which each transit occurred or according to the data subset in which the transit occurred. These parameters were computed and coloring was performed by automatic editing of Snoopy files encoded in xml format, again with the help of a R script.

## Author’s contributions

AP and SP performed single cell time-series experiments and gene expression analyses and evaluated their results. MaHa supervised the experimental work and developed the sample preparation method together with SP. MoHe essentially contributed the analysis of the graph theoretical properties of Petri nets and wrote a corresponding section of the manuscript. WM conceived of and supervised the study, performed computational analyses including the automated generation of the Petri nets and wrote the paper. All authors read and approved the final version of the manuscript.

## Supplementary Tables and Figures

**SI Table 1.** Quantil distributions of the x-fold deviation (x) of a value from the median of values measured in dark controls or far-red stimulated cells. The table quantitatively characterizes the frequency distributions shown in SI Fig. 1B,C.

|  | Dark |  | Far-red |  |
| --- | --- | --- | --- | --- |
| Percent of values | Quantile (Log2(x)) | Quantile (x) | Quantile (Log2(x)) | Quantile (x) |
| 1% | -1.055 | 0.481 | -1.040 | 0.486 |
| 5% | -0.593 | 0.663 | -0.609 | 0.655 |
| 25% | -0.186 | 0.879 | -0.185 | 0.880 |
| 50% | 0.000 | 1.000 | 0.000 | 1.000 |
| 75% | 0.181 | 1.134 | 0.180 | 1.133 |
| 95% | 0.584 | 1.499 | 0.622 | 1.539 |
| 99% | 1.017 | 2.024 | 1.073 | 2.103 |

**SI Table 2.**
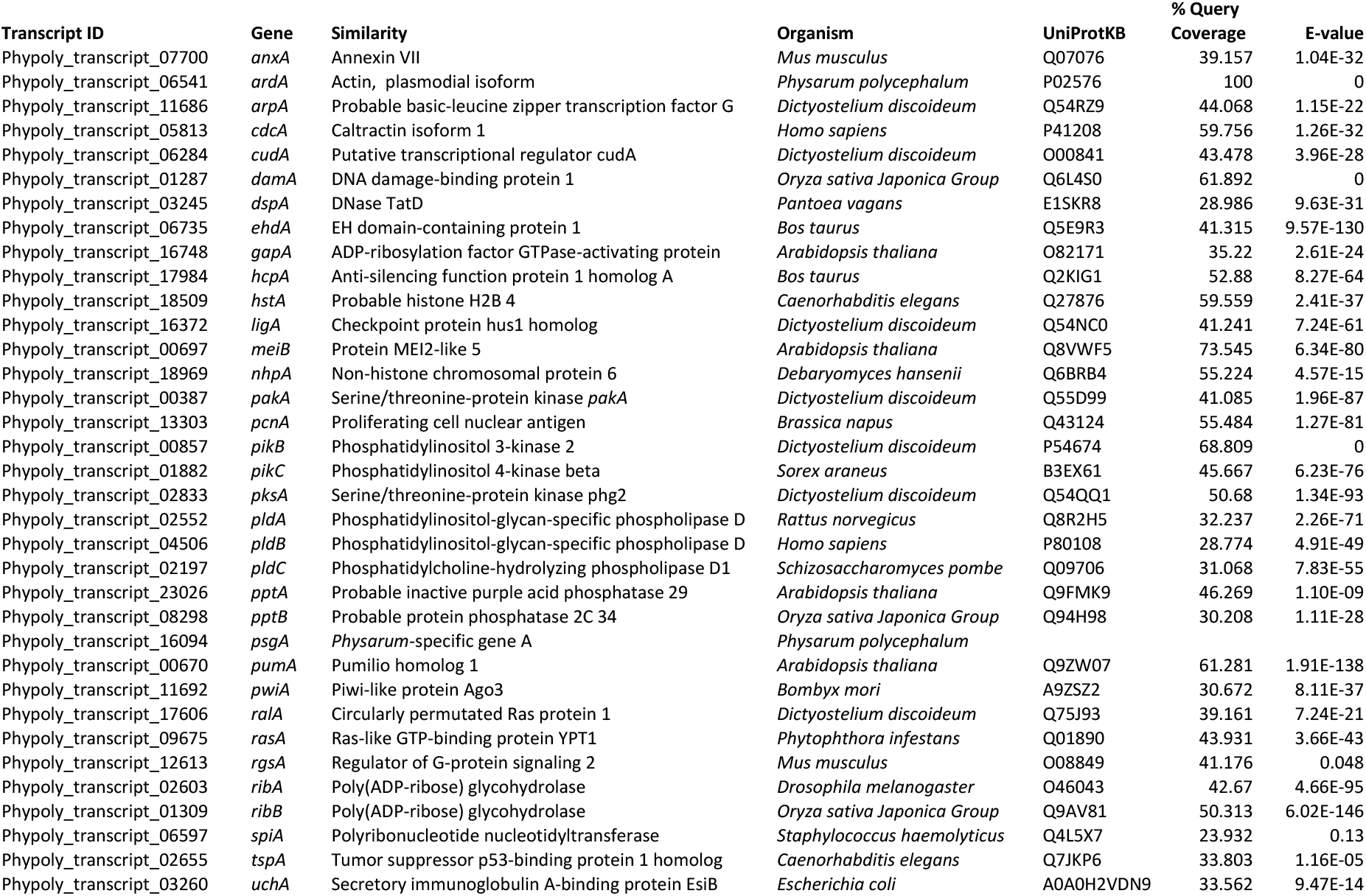
Genes analysed.

**SI Figure 1.**
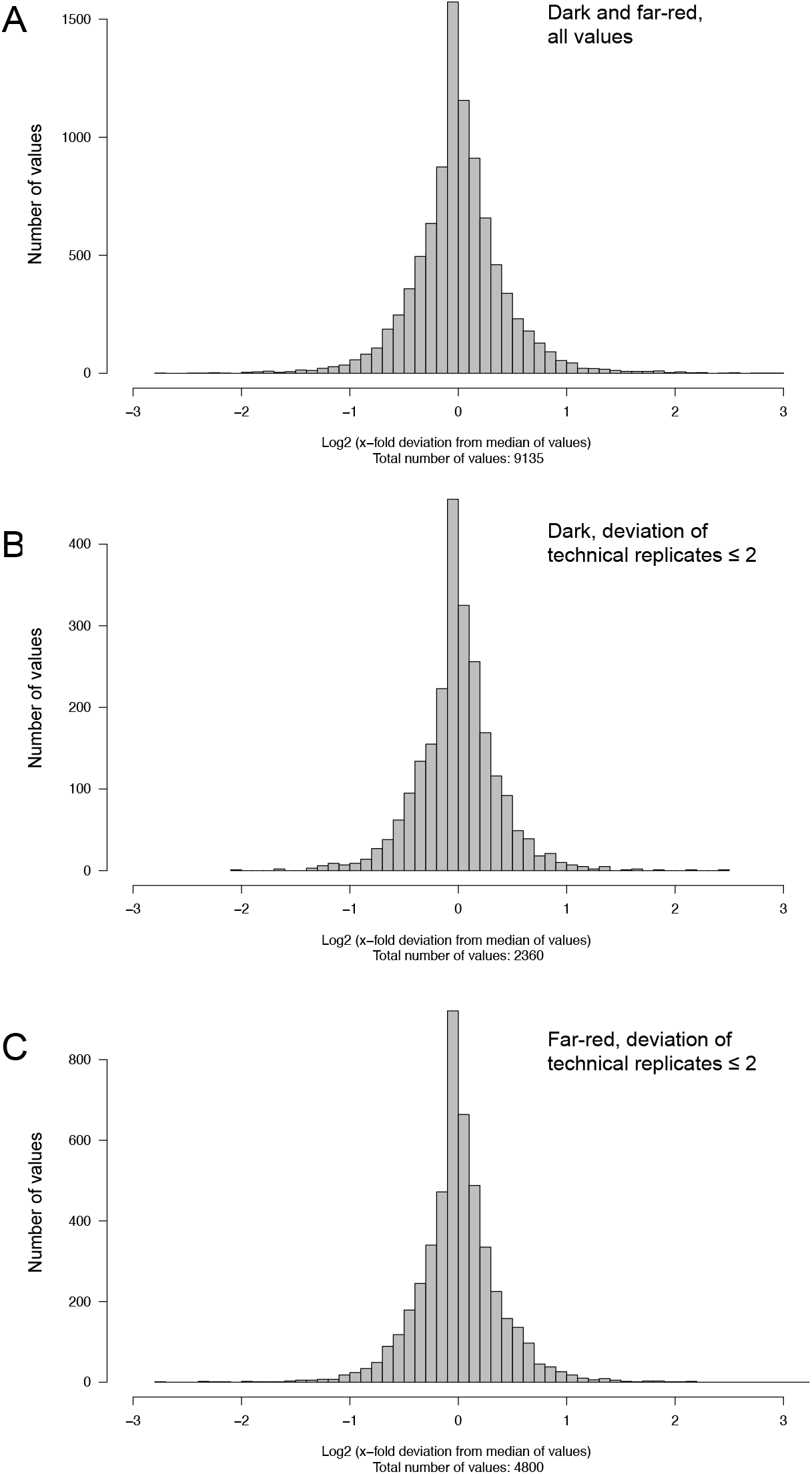
Homogeneity in gene expression, assayed by multiple sampling of individual plasmodial cells. In total, 43 plasmodia (15 dark controls and 28 far-red stimulated plasmodia, analysed at 6h after the stimulus pulse) were evaluated. Nine (3 × 3) or sixteen (4 × 4) samples were simultaneously taken from the same plasmodium and the concentration of the mRNAs of the set of 35 genes (SI Table 2) was determined twice by RT-PCR for each RNA sample. The frequency distributions display the Log2 of the x-fold deviation of each expression value from the median of all values for each gene. Panel (A) displays all values, measured in far-red stimulated cells and dark controls. Panels (B) and (C) display only those measurements where the two values of a RNA sample obtained by technical replication of the RT-PCR differed at maximum by a factor of two. Dark controls (B) and far-red stimulated cells (C) were evaluated separately

**SI Figure 2.**
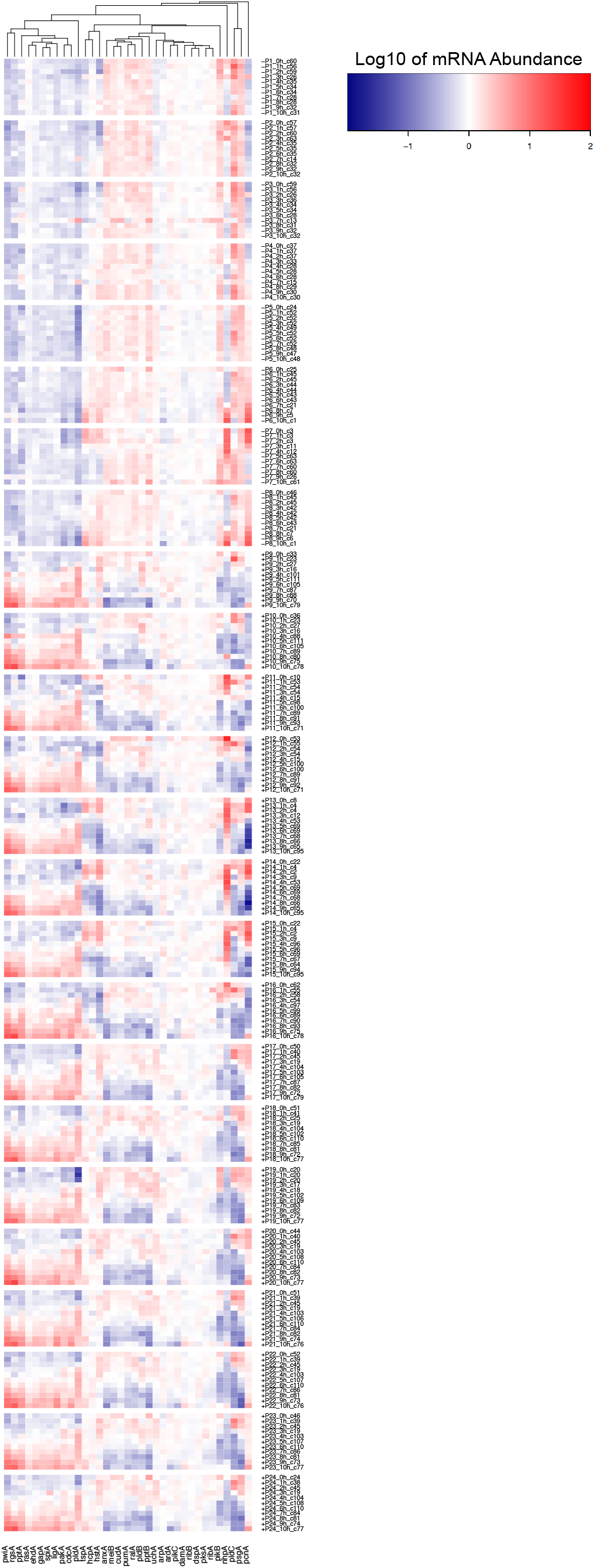
Heat map of time series of gene expression measured in individual plasmodial cells. Plasmodia are numbered according to Table 2. Each line of the heat map represents the gene expression pattern of a cell at the given time point. The label +P12_1h_c55, for example, indicates that the expression pattern of plasmodium number 12 as measured at 1h after the start of the experiment (corresponding to the onset of the far-red light stimulus in light-stimulated cells) was assigned to Simprof cluster number 55 and that the plasmodium had sporulated (+) in response to the stimulus (+, sporulated; -, not sporulated).

**SI Figure 3.**
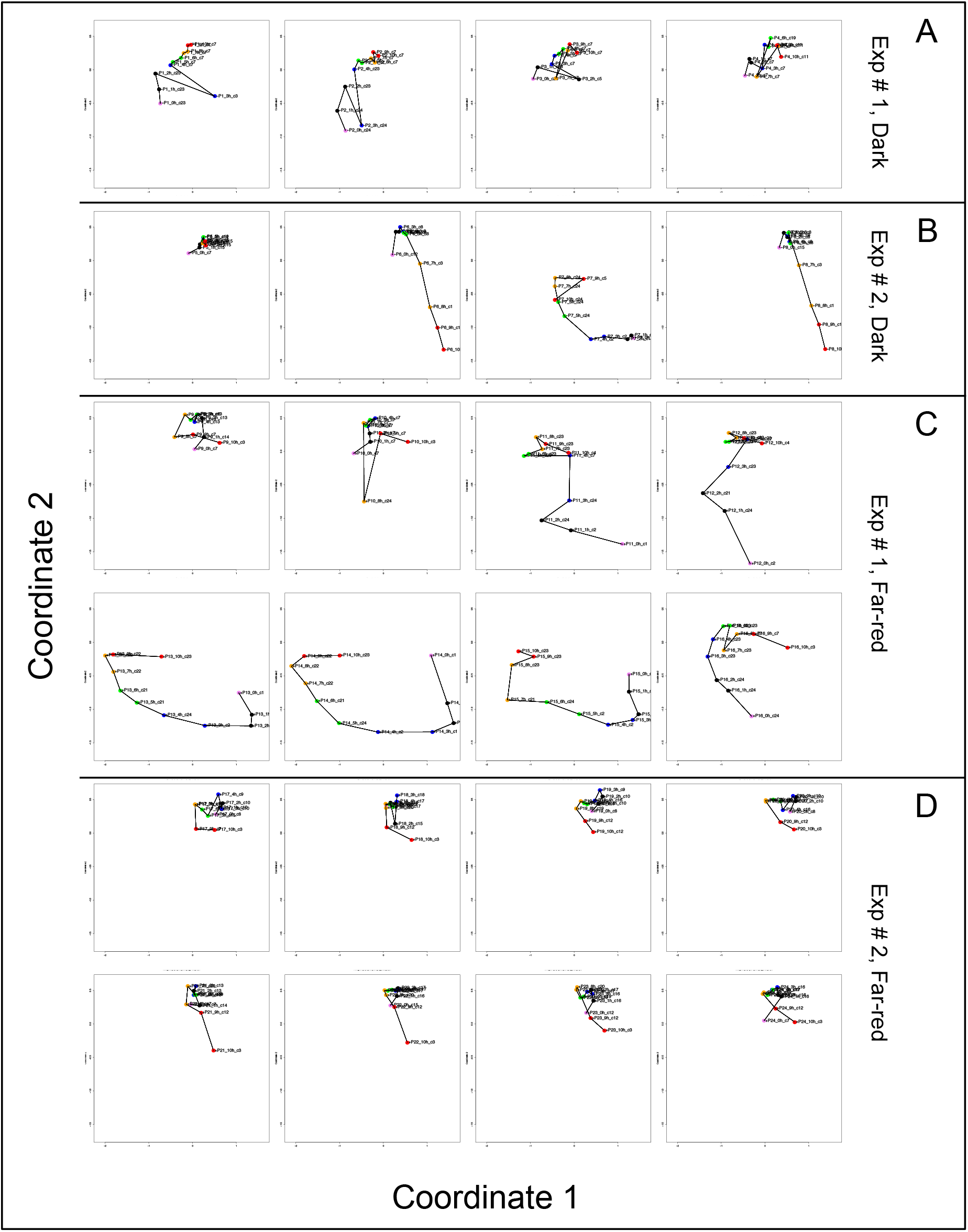
Single cell trajectories of gene expression displayed after multidimensional scaling (MDS) of the expression patterns of the *pcnA* group of genes *(hstA, nhpA, pcnA, uchA)*. For further details see legend to Figure 2.

**SI Figure 4.**
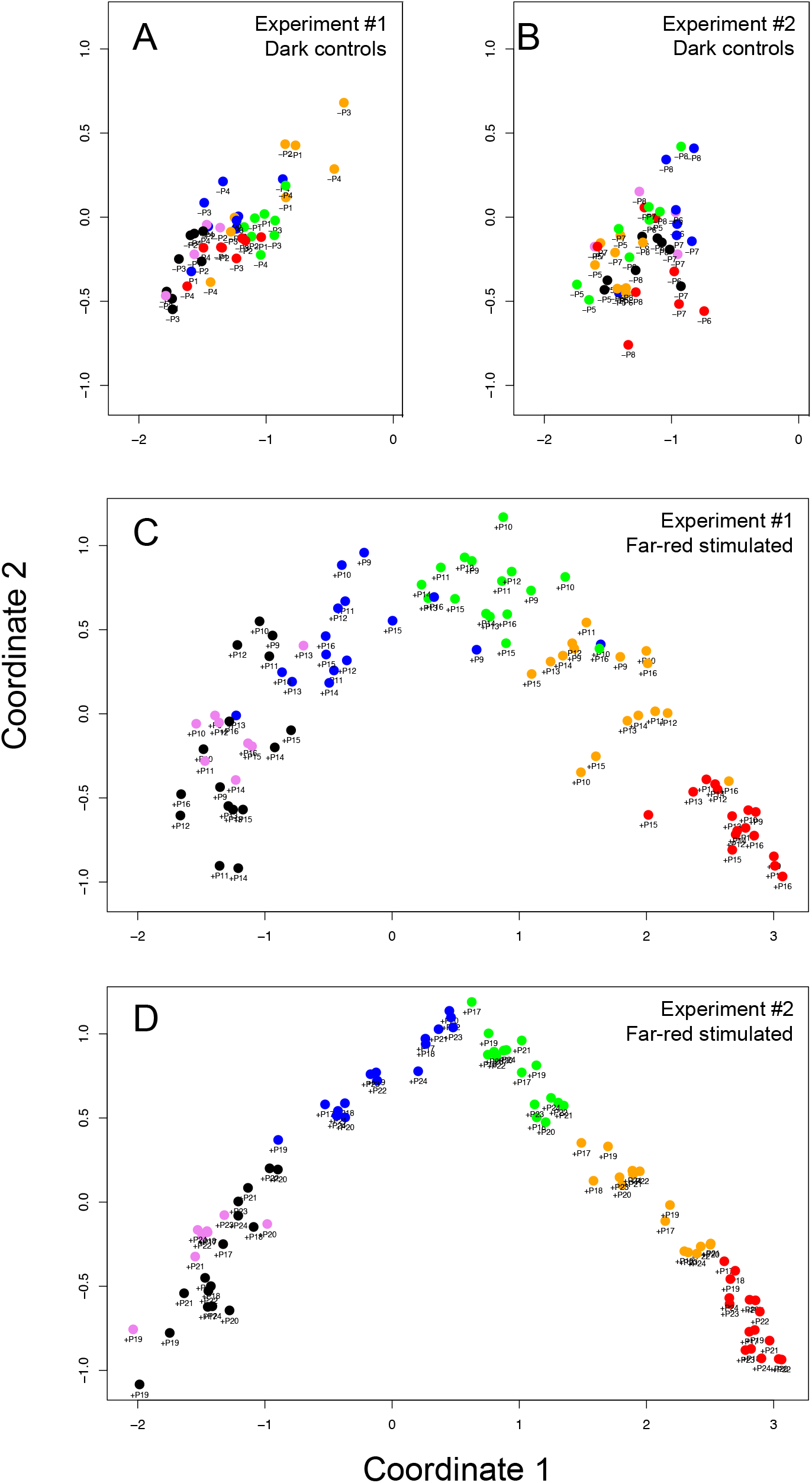
Multidimensional scaling of expression patterns of genes that were obviously up- (*cdcA, ehdA, gapA, ligA, pakA, pldA, pptA, pwiA, rgsA, spiA*) or down-regulated (*anxA, cudA, meiB, pikB, pldB, pldC, pptB, psgA, pumA, ralA*). Time is encoded by color (0h, pink; 1h, 2h, black; 3h, 4h, blue; 5h, 6h, green; 7h, 8h, ocher; 9h, 10h, red).

**SI Figure 5.**
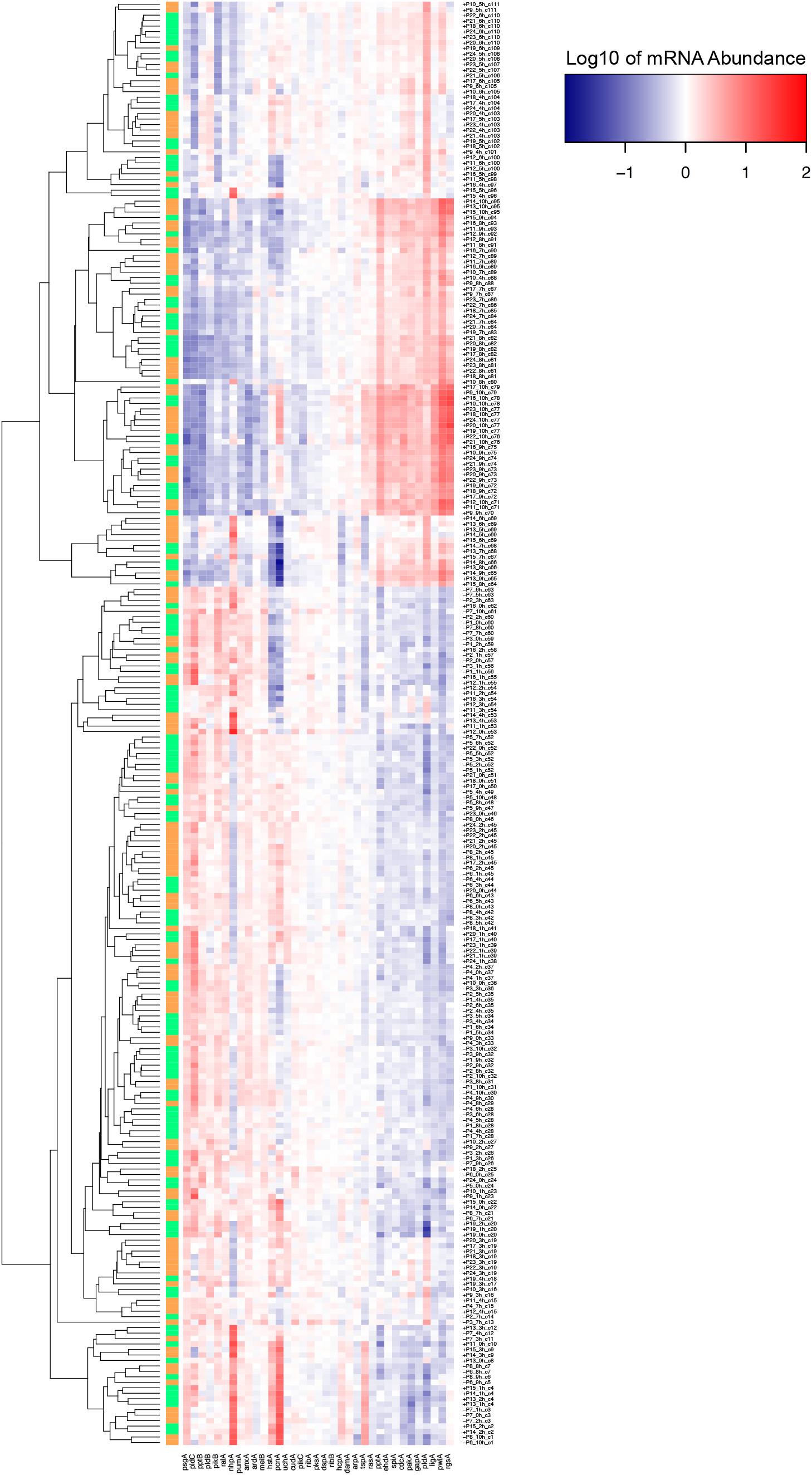
Heat map of gene expression values clustered by the Simprof algorithm. Each line of the heat map represents the gene expression pattern of a cell at a given time point. For labelling of expression patterns see legend to SI Figure 2. The side bar at the left side of the heat map indicates the borders of the individual clusters.

**SI Figure 6.**
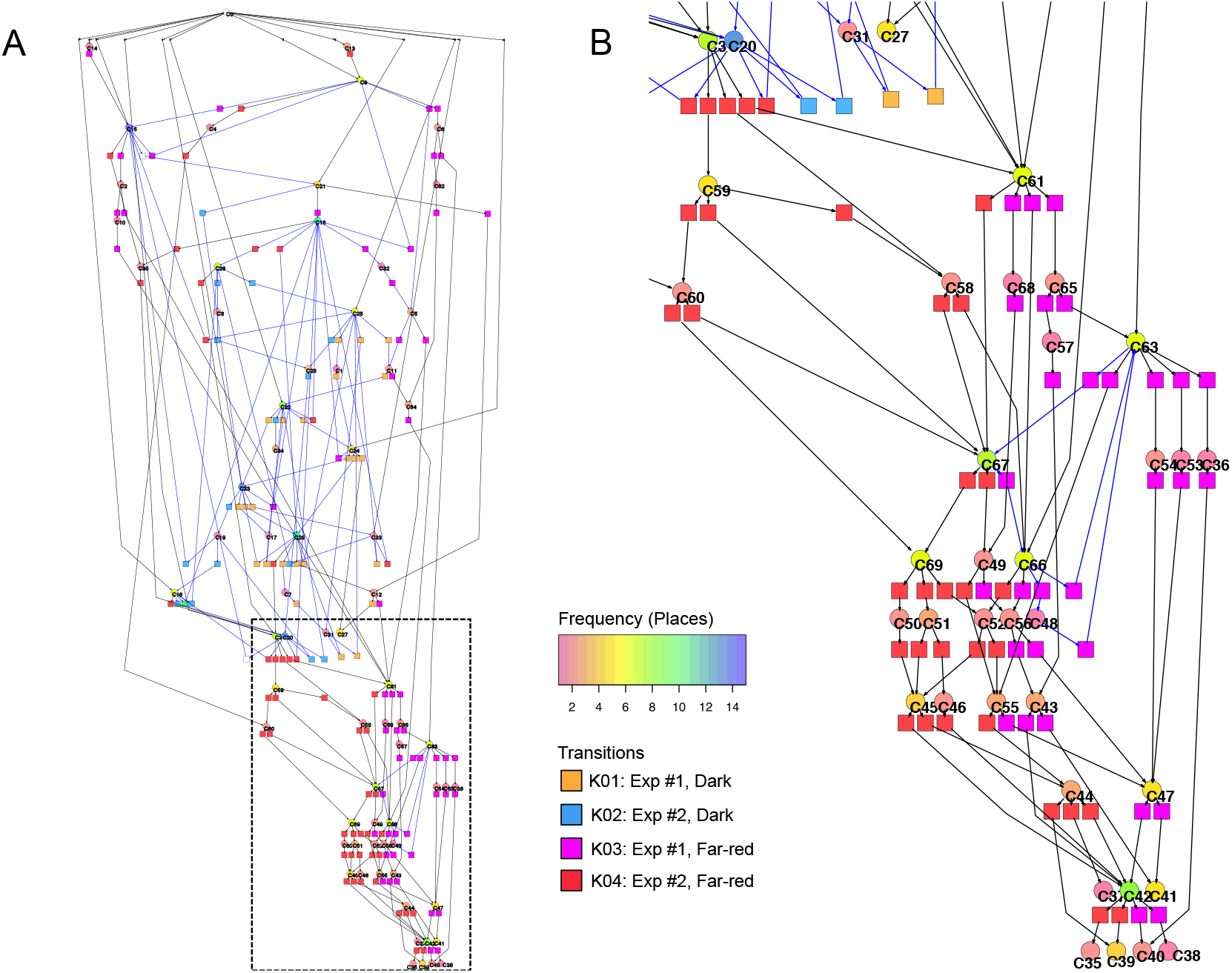
Petri net constructed for the set of up- or down-regulated genes, as listed in the legend to SI Figure 4. (A) Complete Petri net with a high number of T-invariants (indicated by arcs in blue) seen in the upper part of the net. (B) Magnified lower part of the Petri net in panel (A), marked by the rectangle of dashed lines. The part shown in (B) contains mainly transitions corresponding to far-red stimulated cells. Color coding of places indicates the relative frequencies of states of gene expression. Color coding of transitions indicates the group of cells (Table 2) in which the transits occurred.

**SI Figure 7.**
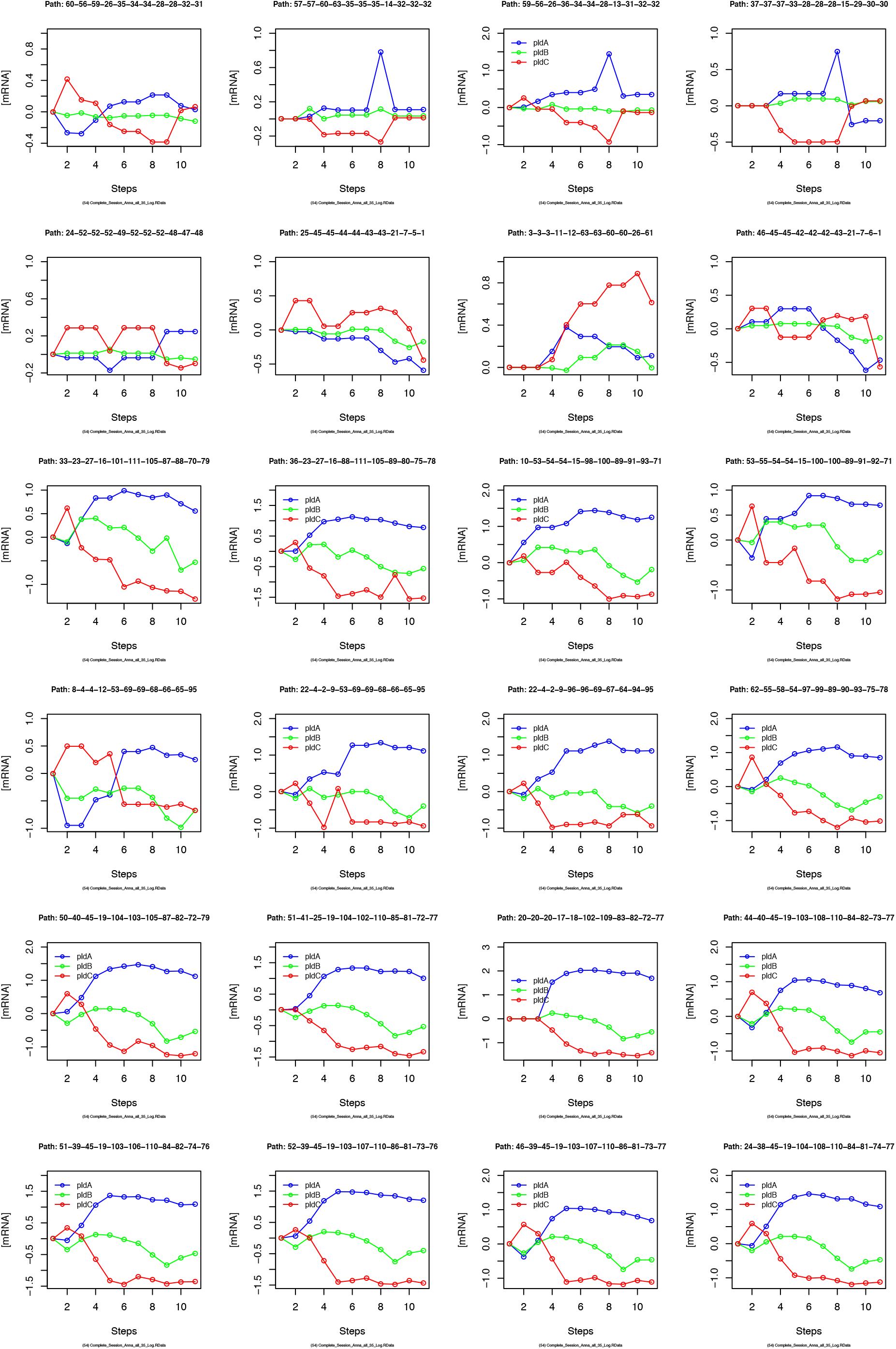
Gene expression kinetics as derived from single cell trajectories. For details see legend to Figure 8.

**SI Figure 8.**
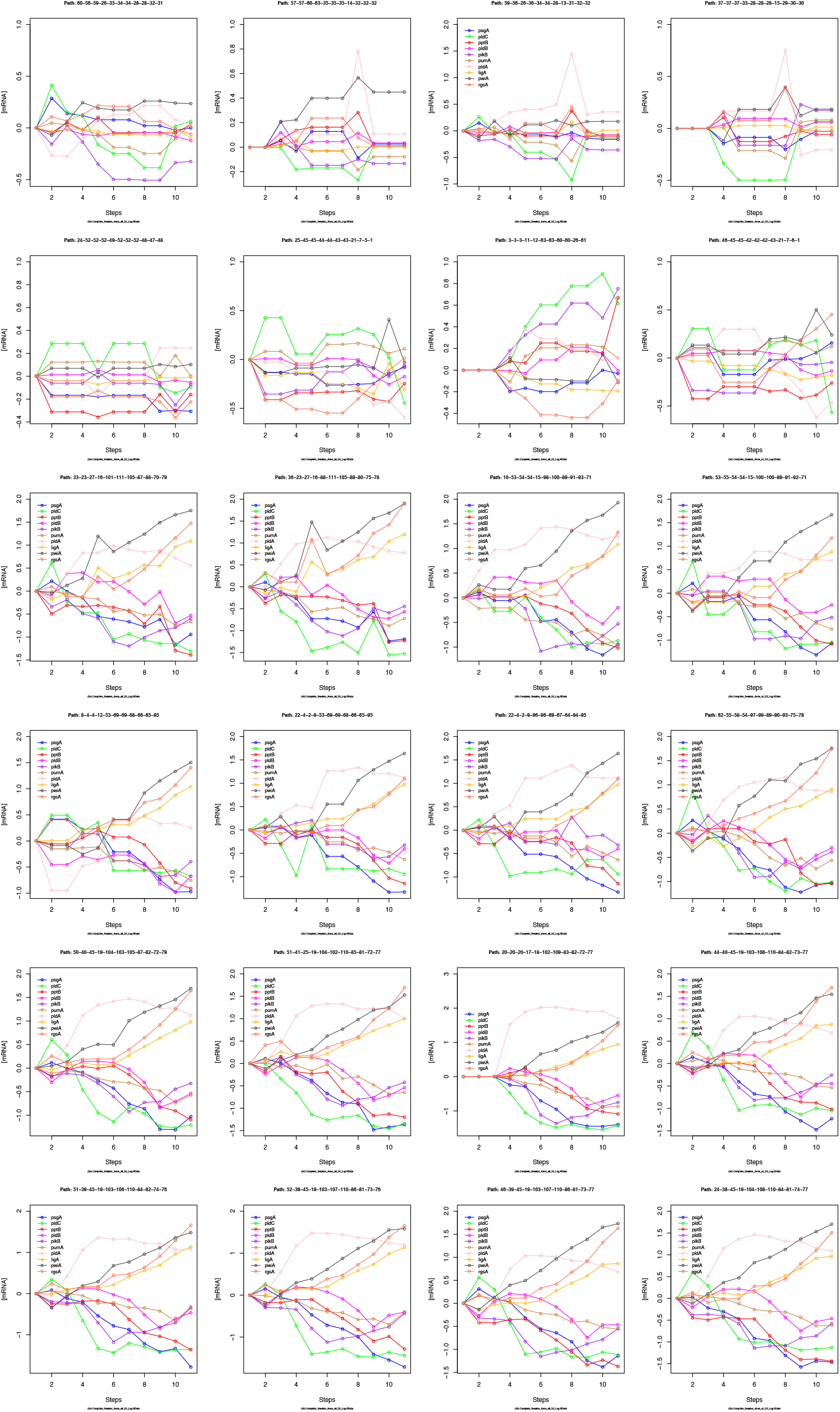
Gene expression kinetics as derived from single cell trajectories. For details see legend to Figure 8

